# Cortical morphology at birth reflects spatio-temporal patterns of gene expression in the fetal human brain

**DOI:** 10.1101/2020.01.28.922849

**Authors:** G. Ball, J. Seidlitz, J. O’Muircheartaigh, R. Dimitrova, D. Fenchel, A. Makropoulos, D. Christiaens, A. Schuh, J. Passerat-Palmbach, J. Hutter, L. Cordero-Grande, E. Hughes, A. Price, J.V. Hajnal, D. Rueckert, E.C. Robinson, A.D. Edwards

## Abstract

Interruption to gestation through preterm birth can significantly impact cortical development and have long-lasting adverse effects on neurodevelopmental outcome. We compared cortical morphology captured by high-resolution, multimodal MRI in n=292 healthy newborn infants (mean age at birth=39.9 weeks) with regional patterns of gene expression in the fetal cortex across gestation (n=156 samples from 16 brains, aged 12 to 37 post-conceptional weeks). We tested the hypothesis that noninvasive measures of cortical structure at birth mirror areal differences in cortical gene expression across gestation, and in a cohort of n=64 preterm infants (mean age at birth = 32.0 weeks), we tested whether cortical alterations observed after preterm birth were associated with altered gene expression in specific developmental cell populations.

Neonatal cortical structure was aligned to differential patterns of cell-specific gene expression in the fetal cortex. Principal component analysis of six measures of cortical morphology and microstructure showed that cortical regions were ordered along a principal axis, with primary cortex clearly separated from heteromodal cortex. This axis was correlated to estimated tissue maturity, indexed by differential expression of genes expressed by progenitor cells and neurons; and engaged in stem cell differentiation, neuron migration and forebrain development. Preterm birth was associated with altered regional MRI metrics and patterns of differential gene expression in glial cell populations.

The spatial patterning of gene expression in the developing cortex was thus mirrored by regional variation in cortical morphology and microstructure at term, and this was disrupted by preterm birth. This work provides a framework to link molecular mechanisms to noninvasive measures of cortical development in early life and highlights novel pathways to injury in neonatal populations at increased risk of neurodevelopmental disorder.

## Introduction

The mammalian cortex is composed of functionally distinct regions organised along broad gradients that reflect spatially ordered and concerted variations of cortical structure and function.[1–6] While the mechanisms behind the emergence of this complex topography are not fully understood, cortical patterning is underwritten by dynamic regulation of gene transcription during gestation.[7,8] During embryonic development, early patterning of the neuroepithelium is established though intrinsic genetic mechanisms [9–12] that regulate early neurodevelopmental processes including neurogenesis and neuronal migrations from around 6 - 8 post-conceptional weeks (pcw) in humans.[11,13] During fetal development, this leads to the establishment and expansion of transient neural structures, including the subventricular zone, preplate, and subplate, and, eventually, formation of the cortex.[11,14,15]

The advent of modern transcriptomic technologies has allowed the precise mapping of cortical gene expression during the human fetal period.[16–18] Gene transcription is highly differentially expressed during prenatal development and varies significantly across cortical areas.[8,16,17,19] Interruption to the precisely timed dynamics of gene transcription during gestation is implicated in the onset of common developmental cognitive and neuropsychiatric disorders.[18,20,21]

Recently, the *postmortem* transcription of thousands of genes across the adult brain has been compiled to form brain-wide, gene expression atlases.[18,22,23] This allows precise comparison between spatial patterns of cortical gene expression and neuroanatomy quantified using Magnetic Resonance Imaging (MRI).[24] Neuroimaging studies have found patterns of gene expression in the adult cortex are mirrored by regional variation in cortical morphometry [25] and functional organisation,[26] and are associated with neuroimaging markers of developmental disorders.[27] Similar databases detailing cerebral gene transcription across the full human lifespan from early embryonic stages to adulthood are now available.[16,18] This has created an unprecedented opportunity to explore the molecular correlates of neuroimaging markers of early brain development.

Advances in neonatal neuroimaging now permit the quantification of developmental neuroanatomy *in vivo* at a higher-resolution than previously possible.[28,29] Imaging studies of the developing human brain shortly after birth have characterised a highly dynamic period of cerebral change defined by significant increases in brain volume,[30,31] cortical thickness and surface area,[31–33] progressive white matter myelination[34,35] and ongoing configuration and consolidation of functional brain networks.[36–41] Several studies have also used diffusion MRI models to study the microstructure of the cortex at around the time of birth, identifying areal patterns of development that may relate to ongoing cellular processes including dendritic arborisation and synaptic formation.[42–44] Further, the truncation of gestation due to preterm birth is associated with widespread alterations in cortical morphometry and microstructure indexed by MRI at the time of normal birth that highlight the sensitivity of noninvasive neuroimaging to detect disruptions in early developmental processes [32,42–49].

The combination of these technologies opens a new window to study early human brain development, facilitating a comparison between patterns of prenatal cortical gene expression and the development of the brain at around the time of birth, as well as providing a platform to test mechanistic hypotheses about the impact of early disruptions to brain development during gestation. The potential of this approach has been previously demonstrated using *post mortem* MRI to reveal a correspondence between genes linked to neural development and the microstructure of the fetal cortex. [50] In this study, we explore the association between *in vivo* measures of cortical morphometry at birth and regional patterns of fetal gene transcription in the human brain. We test the hypothesis that noninvasive markers of neonatal cortical structure mirror areal differences in the timing of cellular processes underlying cortical development, as indexed by differential spatiotemporal patterning of gene expression in the fetal cortex. Additionally, we test whether cortical alterations observed after preterm birth and quantified with MRI are linked with a selective vulnerability of developmental neuronal and glial cell populations in the developing cortex.

We define a principal mode of variation in neonatal cortical structure that is aligned to differential patterns of genes expression in the fetal cortex, enriched for foundational neurodevelopmental processes including neuronal differentiation and migration, and disrupted by preterm birth.

## Results

### A principal axis of the neonatal cortex

Using high-resolution structural and diffusion MRI data acquired from a large cohort of healthy neonates (n=292, 54% male, median [range] gestational age at scan = 40.86 [37.29-44.71]), we extracted six measures of cortical morphology (cortical thickness) and microstructure (T1w/T2w contrast; Fractional Anisotropy, FA; Mean Diffusivity, MD; Intracellular Volume Fraction, fICVF; and Orientation Dispersion Index, ODI) from eleven cortical regions-of-interest (ROI) with corresponding mRNA-sequencing in a prenatal transcriptomic dataset [18] (Figure 1A, Figure S1).

**Figure 1:**
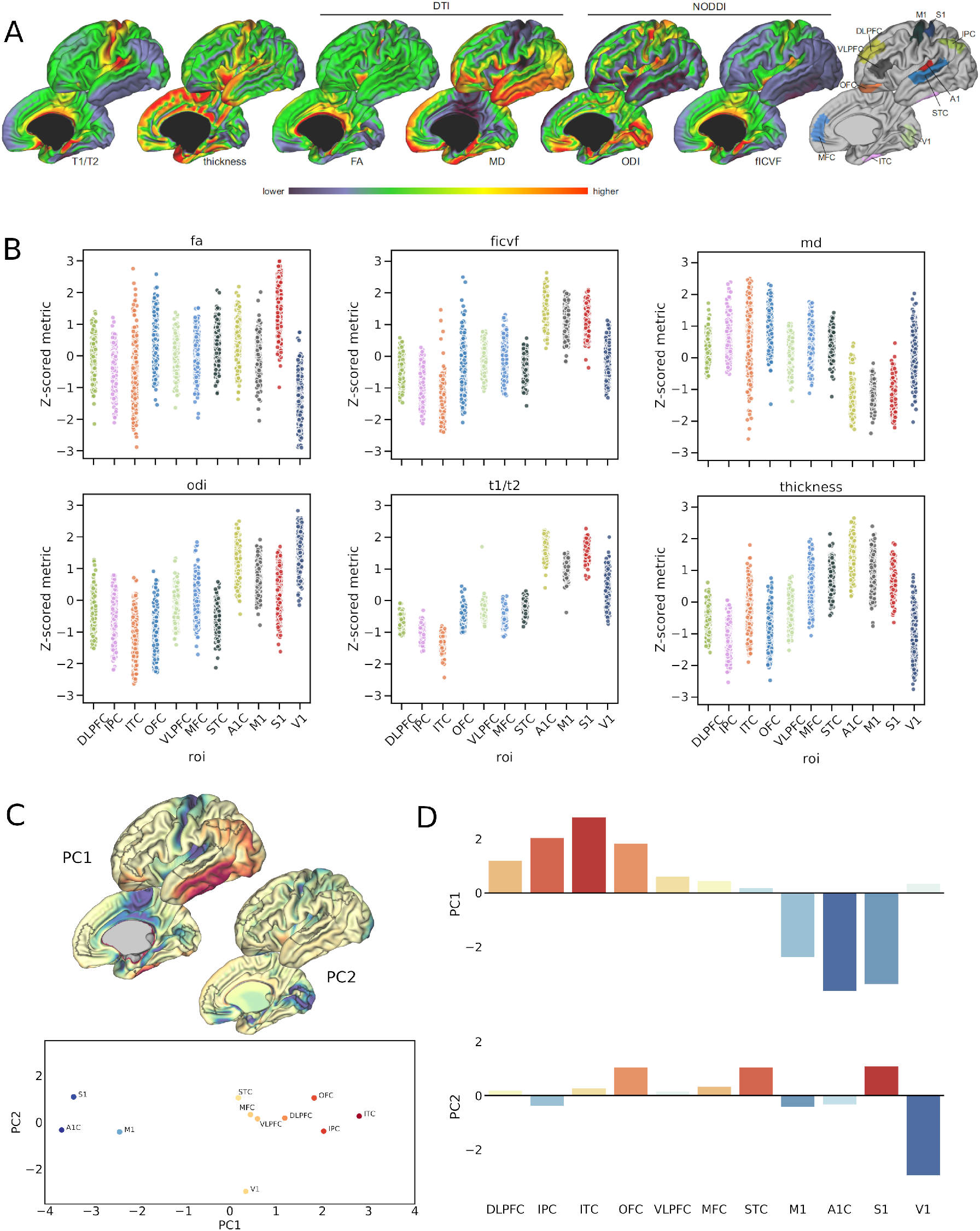
A principal axis of the neonatal cortex indexed by multi-modal MRI. A. Average cortical neuroimaging metrics in a cohort of healthy, term-born neonates (n=292). Metrics derived from structural MRI (T1/T2 contrast, cortical thickness) and diffusion MRI model parameters using DTI (FA and MD) and NODDI (fICVF and ODI). Right. Cortical regions-of-interest based on anatomical references with corresponding developmental transcriptomic data (Figure S1). B. Z-scored cortical metrics are shown for each subject grouped within each cortical region-of-interest. C. Top. Cortical representations of the first two principal components (PC1, PC2) derived from PCA of the regional MRI metric data in B. Bottom. Position of each cortical region-of-interest in PCA state space, the position of each region is dictated by its component score (D) for the first two principal components. Regions are labelled and coloured by PC1 score. D. PCA scores of each metric for the first two principal components, coloured by PC1 score.

Normalised regional metrics for all subjects are shown in Figure 1B. Similar regional profiles are evident across metrics with the pattern of inter-regional variation reflecting the full transcortical patterns shown in Figure 1A. Comparable regional patterns were observed in FA and cortical thickness, and between ODI, fICVF and the T1w/T2w contrast (Figure 1B), with higher FA, thicker cortex and a higher T1w/T2w contrast in primary somatomotor cortex (Figure 1A & 1B). MD displayed an opposing trend across regions, lowest in primary somatomotor regions and highest in fronto-parietal regions.

Based on the similarities in cortical patterning across metrics, we hypothesised that regional variation across metrics could be represented by a small number of latent factors. Using Principal Component Analysis (PCA), we projected the regional metrics onto a set of principal axes that maximally explained variance in the full set of cortical measures (Figure 1C). Using the group-average region × metric matrix, we found that the first two components explained 91.6% of the total variance (PC1 = 72.3% and PC2 = 19.3%, respectively).

The ordering of regions along the principal axis (PC1; Figure 1C) illustrates a clear separation between primary and higher order cortical regions based on neuroimaging metrics, with primary somatosensory and motor cortex (A1C, M1, S1) situated at opposite ends to prefrontal, inferior parietal and temporal cortex (DLPFC, IPC, ITC). This pattern is apparent in all cortical metrics, most strongly in T1/T2 contrast, fICVF and mean diffusivity (Figure S2). The second principal axis (PC2) predominantly captured anatomical and microstructural differences in V1 compared to other primary cortex (Figure 1C & 1D).

### PC1 is associated with regional patterns of gene expression in mid-gestation

Using a developmental transcriptomic dataset of bulk tissue mRNA data sampled from cortical tissue in 16 prenatal human specimens,[18] we compared regional variation in cortical MRI metrics, represented by PC1, with prenatal gene expression in anatomically-correspondent cortical regions. Through comparison to five independent single-cell RNA studies of the developing fetal cortex [18,51–54], we selected a set of 5287 marker genes shown to be differentially expressed in cortical cell populations during gestation. We used a nonlinear mixed-effects approach to model developmental changes in gene expression as a smooth function of age, accounting for inter-specimen variability. The nonlinear model provided a better fit of the expression data for all genes compared to a comparable linear model (range AIC difference: −16.7 to −87.7; range BIC difference: - 2.1 to −58.9).

Using specimen- and age-corrected RPKM values provided by the residuals of the nonlinear mixed model for each gene (Figure S3), we tested the association between spatial variation in gene expression during gestation and regional PC1 score using non-parametric correlation (Kendall’s *τ*). Of 5287 genes, 120 displayed a significant (positive or negative) correlation with PC1 after correction for multiple comparisons with False Discovery Rate (p<0.05). In total, 71 genes were positively correlated with PC1, with increasing gene expression in regions with a higher PC1 score (mean ± S.D. *τ* = 0.208 ± 0.023) and 49 genes displayed the opposite relationship, with higher expression in regions with a negative PC1 score (mean ± S.D. *τ* = −0.208 ± 0.022).

We reasoned that genes associated with the patterning of cortical morphometry at birth may subserve important neurodevelopmental functions. To test this, we performed an over-representation analysis (ORA)[55] for ontological terms associated with specific biological processes in both gene lists. Of 71 genes with spatial patterns of expression positively correlated with PC1 (denoted PC+), 61 (86%) were annotated to specific functional terms. Using all protein-coding genes transcribed in the bulk RNA dataset as the background reference set, we found significant enrichment of several neurodevelopmental terms including: stem cell differentiation (FDR=0.001, enrichment ratio=9.32), neuron migration (FDR=0.03, enrichment=7.94) and forebrain development (FDR=0.004, enrichment=5.65) (Figure 2A; Table S1). Terms relating to stem cell and neuronal differentiation remained significantly enriched when restricting the background reference set to only include genetic markers of fetal cortical cells (n=5287; Table S1). Performing weighted gene correlation network analysis (WGCNA) on the PC+ gene set, we identified two co-expression modules (Figure 2B). The largest, contained 53 genes including a tightly correlated set of developmental genes with roles in regulating cell growth and differentiation including *EOMES*, *NEUROD4*, *SFRP1* and *TFAP2C*. The smaller, second module (Module 2) contained 13 genes, with roles including neuronal signalling (*ERBB4*, *CALB2*, *SCGN*) and neuronal differentiation (*ZNF536*, *DLX1*).

**Figure 2:**
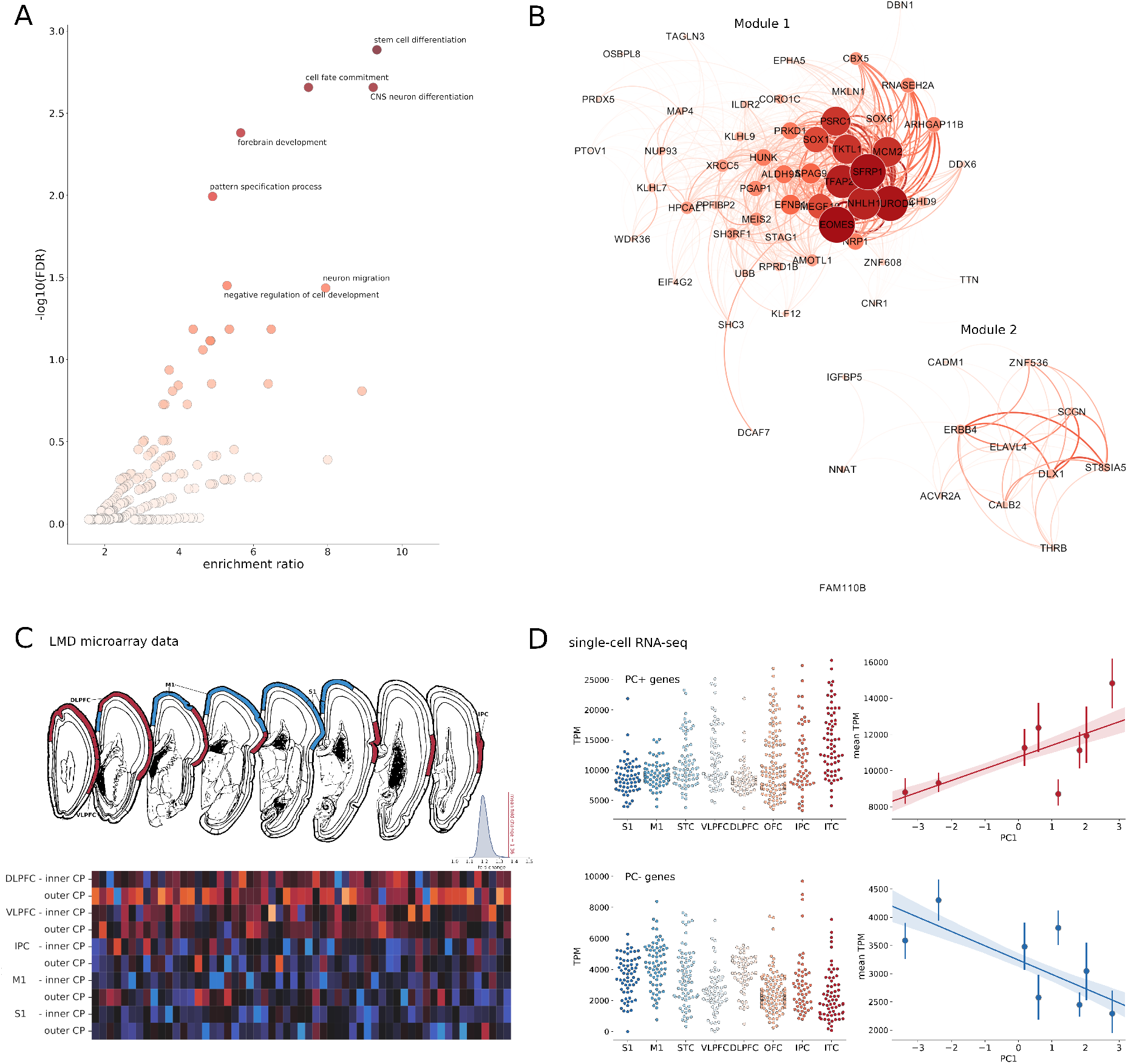
Genes associated with neuronal differentiation are differentially expressed along the principal imaging axis. A. Volcano plot showing enrichment of GO terms (Biological Processes) in genes with age-corrected expression levels positively correlated with PC1. Significantly enriched terms (FDR < 0.05, reference: protein-coding genes) are labelled. B. Gene co-expression analysis of all PC+ genes revealed two modules). Intra-modular connections are shown with node size and colour indicating strength, and edge thickness and colour indicating weight. C. Differential expression of PC+ genes across cortical regions (top) measured using laser microdissection (LMD) microarrays of the cortical plate in two 21pcw fetal samples (https://www.brainspan.org/lcm/). Heatmap shows relative expression of all 71 PC+ genes in the inner and outer cortical plate of each labelled region. Inset: mean fold change of PC+ genes compared to 10,000 random gene sets of the same size. D. Total expression (in transcripts per million, TPM) of PC+ (top) and PC- (bottom) genes in single cells (n=572) extracted from cortical regions in an independent single-cell RNA-seq survey of the mid-gestational fetal cortex [51]. Scatterplots show mean TPM averaged over cells in each region, correlated with each region’s PC1 score.

No biological terms were significantly enriched in genes with a spatial pattern of expression negatively correlated to PC1 (denoted PC-). Using WGCNA, three small modules of 7 genes each were identified (Modules 1N-3N; Figure S4), including genes with high neuronal expression (Module 1N; *CDKL5*, *ZBTB18*, *SORCS1*), and genes involved in cellular processes including adhesion and signalling (Module 2N: *ACTN2*, *PTPN2*, *SSX2IP*) and metabolic activity (Module 3N: *DUSP7*, *ST3GAL1*).

Using independent microarray data from laser microdissections (LMD) of the 21pcw fetal cortex, [16] we verified that PC+ genes had higher expression in the cortical plate of higher order regions (DLPFC, VLPFC, IPC) compared to primary cortex (M1, S1) (Figure 2C; mean fold change = 1.36, p<0.001, 10,000 permutations). Using the top 100 differentially expressed genes identified in the LMD dataset, we also confirmed that genes with higher expression in regions with a higher PC1 score (DLPFC, VLPFC, IPC) in mid-gestation were enriched for important neurodevelopment functions including neuron differentiation (GO:0021953, FDR=0.019, reference: all genes; Table S2). Additional validation experiments using independent single-cell RNA-seq data [51] confirmed an association between regional PC1 score at birth and expression of PC1+ and PC- gene sets in mid-gestation (Figure 2D).

### Imaging-gene associations are enriched for specific cell types in the fetal cortex

To explore these relationships further, we reconstructed cellular gene expression profiles by stratifying the bulk tissue expression data using genetic markers of cell type derived from single-cell RNA studies of the fetal cortex [18,51–54].

Sets of genetic markers for eleven cortical cell classes were initially compiled by combining lists of genes that are differentially expressed in fetal cortical cell populations (Table S2). To verify this grouping, we calculated the average expression trajectories for all genetic markers within each cell type across gestation and used them to calculate a 2D embedding using Uniform Manifold Approximation and Projection (UMAP; Figure 3A). Proximity in the embedded space reflects similarity between average trajectories of gene expression within cell type over time. In the embedded space, cell types clustered by assigned class, and maturational timing (e.g.: precursor or mature), as well as within cellular subtype (eg: inhibitory and excitatory neurons; Figure S5).

**Figure 3:**
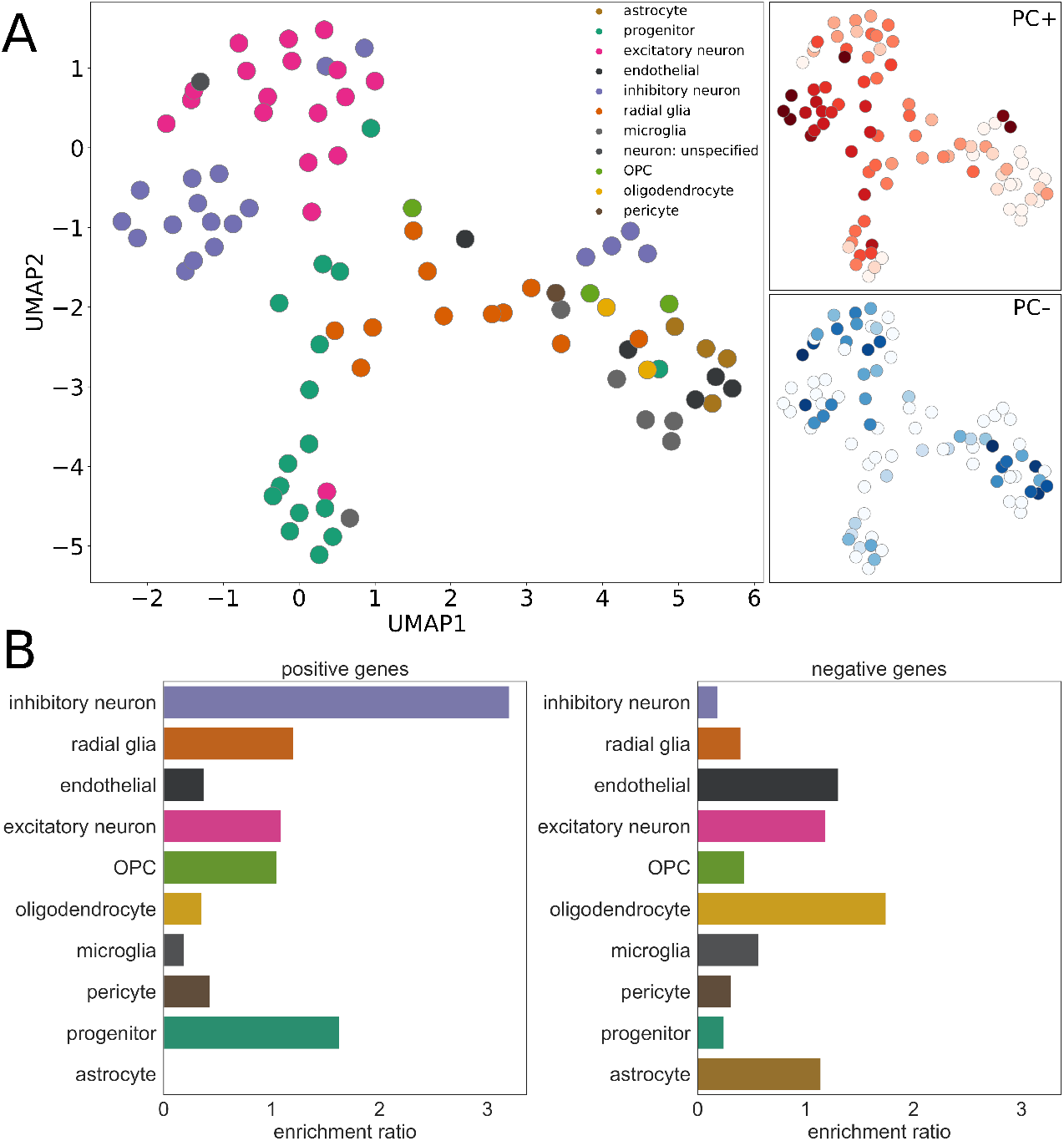
Cell-specific gene expression is associated with cortical morphology at birth. A. UMAP embedding of 86 cell types based on trajectories of relative gene expression over time recovers annotated cell classes. Subplots reflect enrichment ratios of cell classes in PC+ and PC- gene sets (darker colour represent higher enrichment ratio). B. Enrichment ratio for fetal marker genes expressed by each cell class is shown for PC+ (left) and PC- (right) gene sets.

We tested the enrichment of genes expressed by each cell class within the PC+ and PC- gene sets. We found that PC+ genes were significantly enriched for genes expressed by precursor cells (p=0.0003, reference: fetal gene markers), specifically, for genes expressed by intermediate progenitor cells (enrichment ratio = 1.63, p=0.0002; Table S4), and inhibitory neurons (enrichment ratio = 3.2, p<0.0001; Figure 3B; Table S4). Posthoc analysis within cell class revealed specific inhibitory neuron subtypes present in the mid-fetal brain and enriched in the PC+ gene set included migrating cortical interneurons from the caudal ganglionic eminence (In_5,[51] IN-CTX-CGE2,[52] both p<0.0001) and newborn interneurons originating in the medial ganglionic eminence (nIN1;[52] p=0.0017).

In contrast, PC- genes, with a spatial pattern of expression that was higher in primary somatomotor regions at mid-gestation, were enriched for genes expressed by mature cell types (enrichment=1.18, p=0.002). In terms of cell class, genes expressed by oligodendrocytes were enriched within PC-, though not significantly (enrichment=1.75, p=0.056; Figure 3B; Table S4). When considering only marker genes uniquely expressed by each cell class, PC- genes were enriched for excitatory neuronal genes (enrichment=2.14, p=0.008; Table S5). Posthoc analysis within this class revealed a single enriched early maturing excitatory neuronal subtype (Ex_4,[51], p<0.0001). Similar patterns of cell class-specific expression of PC+ and PC- genes were observed in the single-cell RNA dataset (Figure S6).

### Variation in tissue maturation during gestation predicts cortical development at birth

These data suggest that the spatial patterning of gene expression in the developing cortex is mirrored by regional variation in cortical morphology and microstructure measured using MRI at birth. To test this hypothesis, we created a model of cortical maturity to capture the relationship between the regional timing of gene expression and tissue maturation.

We modelled tissue sample age as a function of gene expression using support vector regression. To ensure full coverage across the prenatal period, and to maximise the number of samples contributing to the model, we included additional data from all tissue samples from brains aged 8pcw to 4 months postnatal age (n=23 total). Using mean cortical gene expression of all 120 (PC+ and PC-) genes (Figure 4A) our model accurately predicted sample age across the full prenatal window (Figure 4B), up to 4 months of age. We validated our model in a separate dataset comprising microarray data from the prefrontal cortex in n=46 brains aged 13pcw to 4 months [56] (BrainCloud; Figure 4B).

**Figure 4:**
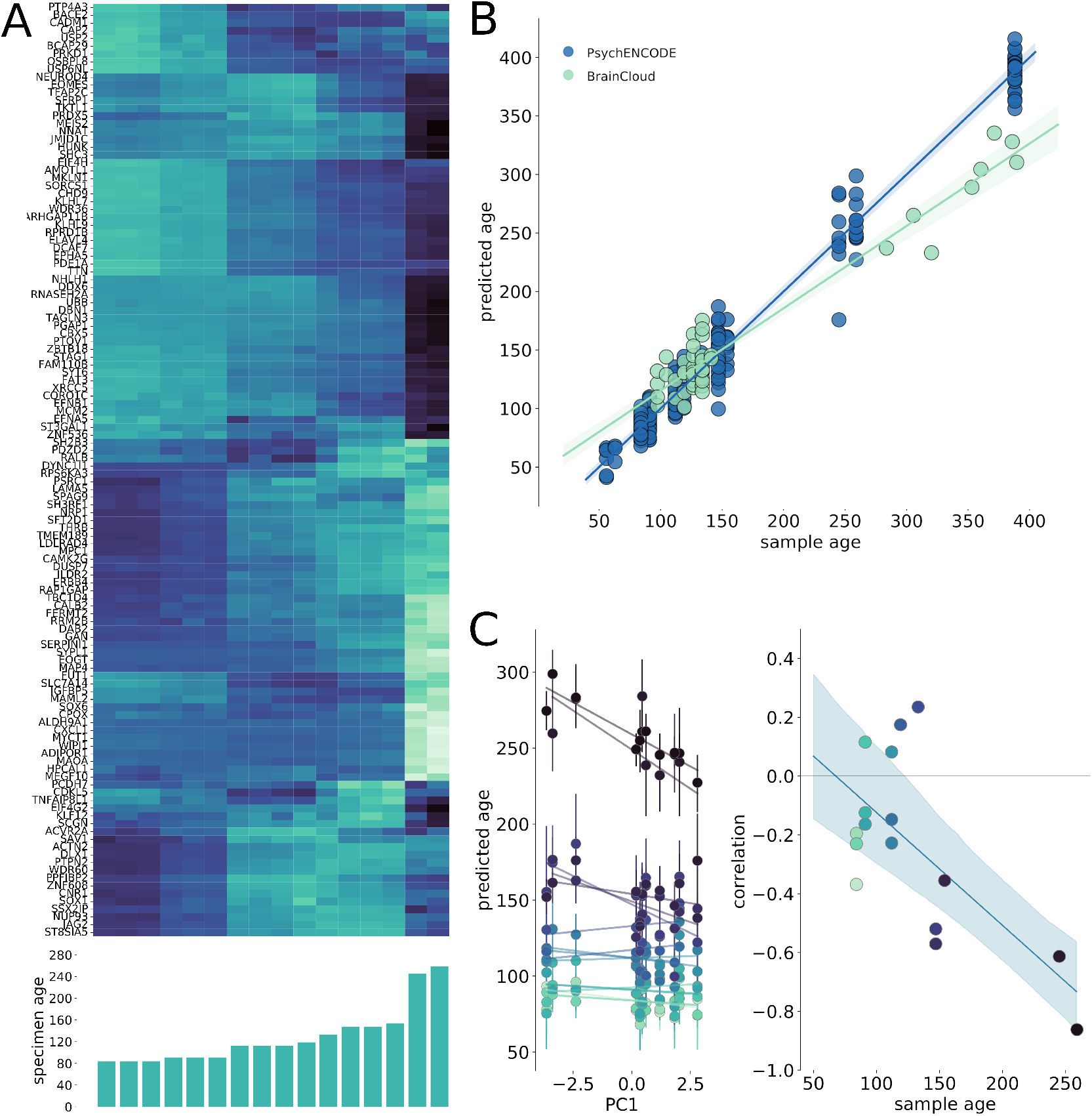
Tissue maturity correlates with regional variation in cortical morphometry at birth. A. Developmental patterns of mean cortical gene expression illustrated in each specimen for all 120 regionally variant genes (PC+ and PC-), ordered by age. B. The relationship between predicted and true sample age for all regional samples (n=198 samples from n=23 brains) in the PsychENCODE dataset aged between 50 and 400 postconceptional days (8pcw to 4 postnatal months), estimated using support vector regression (SVR) and leave-one-out cross-validation. The SVR model was validated using additional samples from the BrainCloud dataset (n=46 samples). Shaded area indicates 95% C.I. C. Left. The correlation between regional PC1 score and predicted tissue maturity is shown for each sample during gestation. Error bars show 95% C.I. for regional age predictions over 1000 bootstrapped gene samples. Right. PC1 correlation is plotted against specimen age for each brain. Shaded area indicates 95% C.I. for linear model fit over bootstrap samples.

Using predictions from this model, we estimated the correlation between regional age predictions and PC1 in the prenatal sample (Figure 4C, left). We expected that for a given brain, regions with a more advanced gene expression profile (i.e: more similar to older tissue samples) would return an older age prediction. We observed a negative association develop over gestation between a cortical region’s predicted maturity and its position along the principal axis at birth (Figure 4C, right; R^2^=0.36, p<0.001 [5000 permutations]), such that in older samples, a lower PC1 score was associated with an older predicted age based on gene expression.

Using nonlinear models of gene expression over time, we estimated regional genetic maturity at several points across gestation (Figure S7). We found that the relative maturity of regions compared to the rest of the cortex varied over time. Primary somatomotor regions remained relatively advanced throughout gestation compared to the rest of the cortex. In contrast, V1 remained relatively delayed across gestation. A divergence in maturity becomes apparent within higher-order regions by mid-gestation, with some cortical areas (IPC, ITC) falling behind other regions towards the time of birth. These patterns were largely repeated using the full fetal gene marker set (n=5287 genes, Figure S8).

### Preterm birth leads to alterations along the principal imaging axis

Based on this evidence, we hypothesised that an interruption to the length of gestation would yield differences in cortical morphology indexed by variation along PC1. To test this, we compared cortical morphology in healthy neonates (n=292) to a cohort of preterm-born infants scanned at term-equivalent age (n=64, 59% male; mean [S.D] gestational age at birth = 32.00 [3.88] weeks).

We extracted neuroimaging metrics from each cortical region and projected each individual’s region × metric matrix onto the principal imaging axis (Figure S9). After correcting for age at scan and sex, regional variation along PC1 explained significantly less variance in preterm individual’s imaging data than those born at term (ANCOVA: F=7.9, p=0.005; Figure 5A). Across both groups, the mean variance explained by PC1 increased with age (Figure 5A; F=46.0, p<0.001), with a stronger association in the preterm cohort (interaction: F=6.63, p=0.01) suggesting that arrangement along the principal axis is ongoing around the time of birth and altered by events surrounding preterm birth. There was no significant difference between sexes (F=1.12, p=0.28).

**Figure 5:**
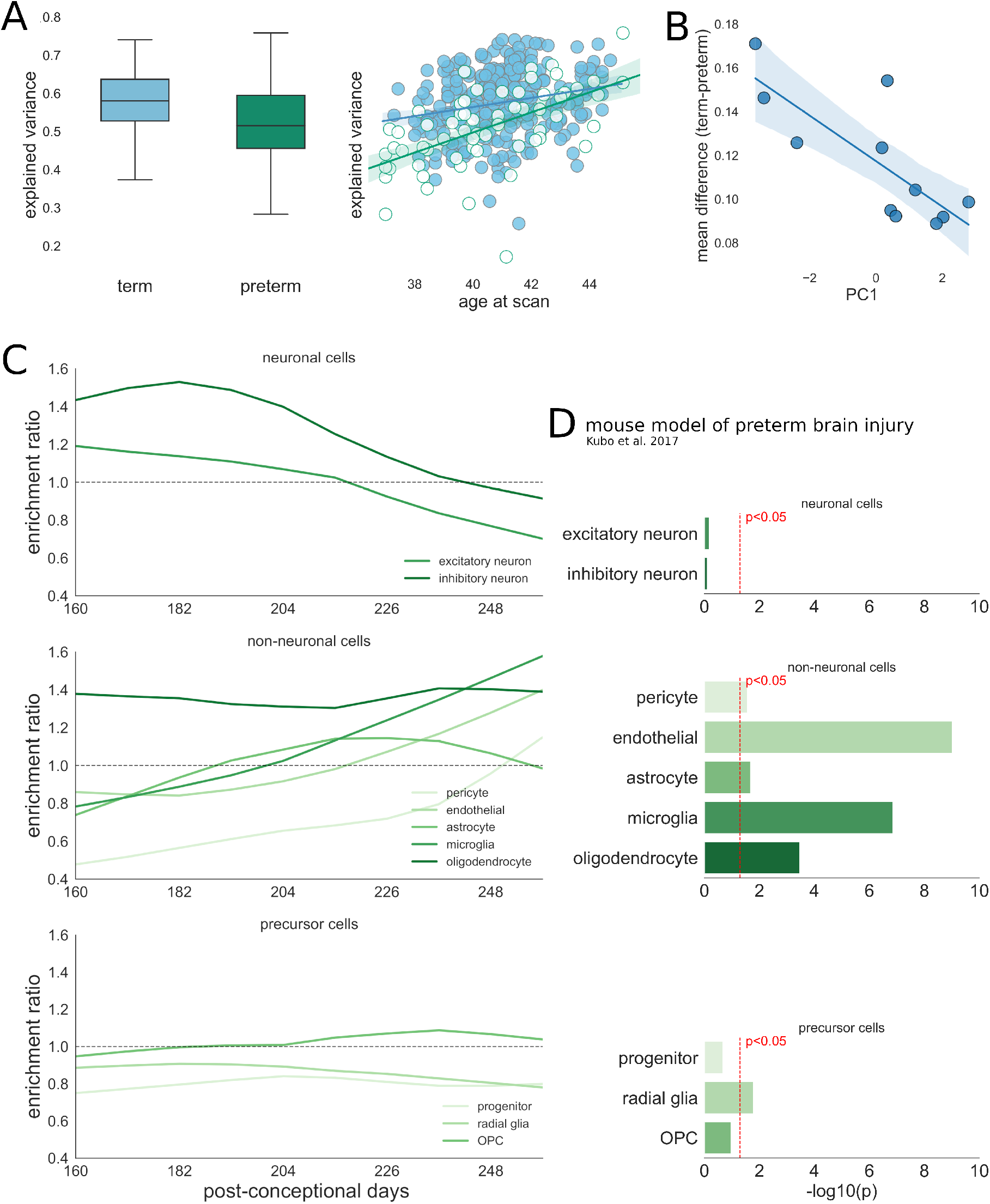
Disruption of the cortical development in preterm-born infants. A. Left. Group difference in individual variance across multiple neuroimaging metrics explained by the principal imaging axis in term (blue) and preterm (green) infants (left). Right. The relationship between age at scan and variance explained by PC1 across all cortical metrics (right). Regression lines are shown for term (blue) and preterm (green) infants with 95% C.I. B. Group differences in regional T1w/T2w contrast ordered by position along PC1. (linear regression shown with 95% C.I.). C. Enrichment of gene sets from 10 fetal cortical cell classes (top: neuronal; middle: non-neuronal; bottom: precursor) based on genes significantly associated (FDR p<0.05) with group differences in T1w/T2w contrast at 10 timepoints in the preterm period. D. Enrichment of genes both expressed by each cell type and significantly correlated with T1/T2 in at least one age window in differentially-expressed genes measured in an experimental mouse model of preterm brain injury. [57]

As differences in the variance explained by PC1 are dictated by individual differences in cortical metrics, we sought to test the specific effects of preterm birth on all imaging measures. Using mixed effects linear models including effects of age, birth status and regional PC1 score, we confirmed a significant main effect of birth status on all cortical metrics except for ODI (Table S6-8).The largest effect was evident in cortical T1w/T2w contrast (F_1,354_=135.53, p<0.0001, Cohen’s *d*=1.62; Table S8). On average, cortical T1w/T2w was significantly lower in preterm infants (marginal means [95% C.I.] = 1.32 [1.31,1.33], 1.20 [1.18,1.22] for term and preterm infants, respectively). To a lesser extent, both intracellular volume fraction and FA were, on average, higher in term infants (d=0.32, 0.56 respectively), although the direction of this effect was not consistent across cortical regions (Fig S10). In contrast, average cortical mean diffusivity (*d*=-1.17) and, to a lesser extent, cortical thickness (*d*=- 0.65) were higher in preterm infants across all regions.

The magnitude of regional group differences across all cortical metrics varied as a function of PC1 (Table S7; Figure 5B; Figure S10). This effect was most apparent in T1w/T2w contrast where the differences between term and preterm groups formed a strong negative association with PC1 (r = - 0.78, p=0.023 after FDR correction). Similar trends were seen in the other metrics, although none reached significance (|r| = 0.32 to 0.68, all p>0.05).

### Vulnerability of specific cell populations to the timing of preterm birth

These data show that cortical differences in preterm infants occur along the principal imaging axis and are most apparent in T1/T2w contrast. We investigated the potential that the differences observed in preterm cortex may reflect a selective vulnerability in specific cell populations due to coincidental timing of extrauterine exposure following preterm birth and temporal variations in gene expression. Focusing on the cortical differences observed in T1/T2w contrast, we first estimated gene expression trajectories over the latter stages of gestation (160 to 260 post-conceptional days, approximately 25 to 39 weeks gestational weeks). We then split this period into 10 age windows and within each, we identified genes with expression significantly correlated to the magnitude of group differences in T1/T2w contrast at term-equivalent age (FDR-corrected p<0.05, Fig 5C). Within each window, we tested for enrichment of gene expression by each of 10 fetal cell types. In the early preterm period, we found that mean regional differences in T1w/T2w contrast at term-equivalent age were significantly associated with genes expressed by both inhibitory and excitatory neurons (windows 1, 2, 3 and 5, hypergeometric statistic: p<0.05; Figure 5C, top; reference: fetal cell markers). However, later in gestation, T1w/T2w differences were correlated with the expression of genes enriched for glial cell populations, including microglia, endothelial cells (windows 8,9 and 10; all p<0.05) and oligodendrocytes (windows 1,2 and 8,9 and 10, all p<0.05; Fig 5C, middle).

Using gene expression data from a mouse model of preterm brain injury [57], we confirmed the relationship between preterm brain injury and altered gene expression in glial cell populations at birth (Figure 5D). Gene expression in the murine cortex was measured at P1.5, after hypoxic-ischemic insult at E16.5. Differentially expressed genes (DEG; p<0.05) were mapped to human homologs with 217 DEGs matched to human genes included in the current study. We found that the set of DEGs was enriched for genes both expressed by glial populations in the human fetal cortex and associated with T1/T2 differences in the neonatal cortex (Figure 5D; Table S9). Relative to a background set of mapped genes (n=15,052) we observed a significant enrichment of genes associated with T1/T2 in at least one age window and expressed by glial populations including microglia (enrichment ratio=6.9, p<0.001), endothelial cells (5.4, p<0.001), oligodendrocytes (4.9, p<0.001) and radial glia (2.2, p=0.02). These relationships remained significant in microglia and endothelial cells when restricting the background set to only include matched fetal gene markers (n=4733).

### Potential cellular processes disrupted in the preterm brain

To identify potential molecular pathways associated with the neuroimaging differences we observed in the preterm cortex, we identified genes expressed by glial cell types associated with both neuroimaging differences in the human preterm brain and experimental models of preterm brain injury. Using genes expressed by oligodendrocytes (Figure 6A), microglia (Figure 6B) and endothelial cells (Figure S11) and associated with T1/T2 differences across multiple prenatal age windows, we identified protein-protein interaction networks using the STRING database [58]. Networks for oligodendrocytes and microglia are shown in Figure 6. We performed a functional enrichment analysis of Reactome pathways [59] using the whole genome as a reference to identify specific molecular processes involving genes in each PPI network and identified significantly enriched pathways in each cell population (Table S10-S12).

**Figure 6:**
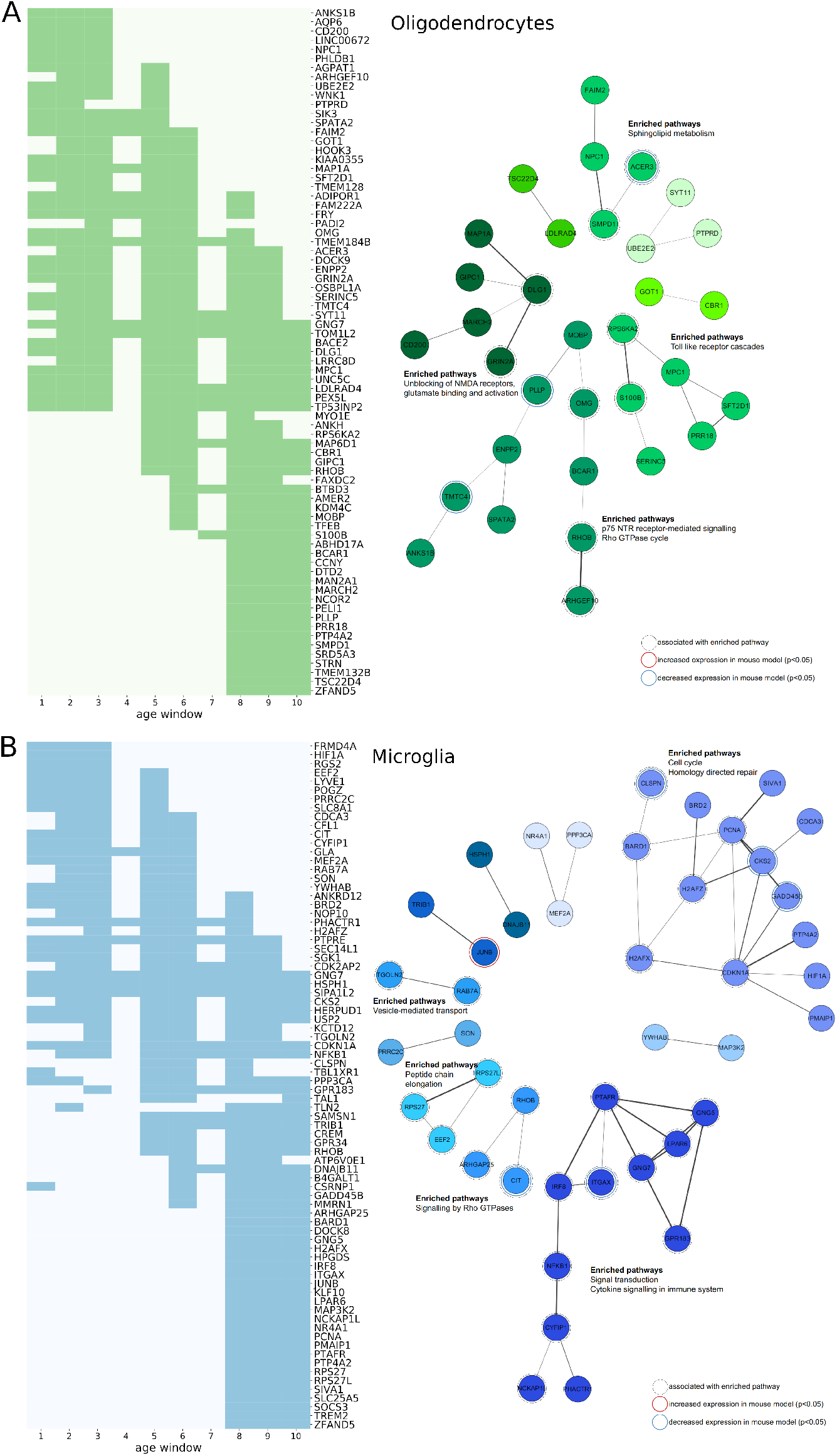
Cellular pathways associated with developmental alterations in the preterm cortex. A. Left. Genes expressed by oligodendrocytes in the fetal cortex and significantly associated with group differences in T1w/T2w contrasts across at least 3 age windows are shown. Dark green indicates periods where gene expression and T1w/T2w contrast were significantly correlated for each gene (FDR p<0.05) across the preterm period. Right. Protein-protein interaction networks derived using STRING. Top functional enrichments of molecular pathways are shown where applicable. Genes associated with listed enriched pathway and genes differentially-expressed in an animal model of preterm brain injury are highlighted. B. Genes expressed by microglia in the fetal cortex. Details as in A.

In oligodendrocytes, pathway enrichment analysis revealed significant gene associations across multiple time windows. Genes involved in NMDA signalling in the MAPK/ERK pathway (HSA-438066, HSA-442729, HSA-442982; *DLG1, GRIN2A*) were significantly correlated to T1w/T2w differences across the majority of the preterm period. In contrast, regional expression of genes associated with the MyD88 and TLR signalling cascades (HSA-975871; *S100B*, *RPS6KA2*) were most closely correlated to T1w/T2w differences in the latter stages of gestation (windows 5 to 9 and 7 to 10, respectively). Other pathways linked genes expression over multiple time periods. Neurotrophin signalling pathways included genes *OMG* (correlated between windows 1-8) and *ARHGEF10* (windows 2-5), and the Rho-GTPase signalling pathway (HSA-194840) included both *ARHGEF10* and *RHOB* (windows 5-10). Finally, sphingolipid metabolism pathways included genes expressed across both the full prenatal in humans and differentially-expressed after fetal ischaemic insult in mice (*ACER3*, windows 2-9).

In microglia, enriched pathways among genes associated with T1/T2 differences in the preterm cortex included signal transduction (HSA-162582), cytokine signalling (HSA-1280215) and stress response (Homology directed repair; HSA-5693538). The regional expression of genes associated with Rho GTPase signalling (HSA-194315; *CIT*, *RHOB*, *ARHGAP25*) spanned the prenatal period and were linked to brain injury in the mouse model (*CIT*). Similarly, *ITGAX*, associated with T1/T2 differences late in gestation and linked to cellular inflammatory responses, was differentially expressed in the mouse brain after fetal hypoxic-ischaemia.

In endothelial cells, a large PPI network enriched for genes associated with apoptotic mechanisms, including *CASP3*, *CDKN1A*, and *GADD45B,* each with patterns of expression correlated with T1/T2 differences in the preterm cortex (Figure S11).

This highlights metabolic signalling pathways associated with genes expressed in developmental glial populations during the period most at risk of interruption by preterm birth with a regional specificity correlated to neuroimaging markers of preterm brain injury at birth and a functional role in experimental models of preterm brain injury.

## Discussion

In this study, we aimed to test the hypothesis that noninvasive neuroimaging measures of cortical structure at birth encode differential spatiotemporal patterning of genes underlying corticogenesis. We found that gene expression in the fetal cortex is mirrored by a principal mode of variation across multiple MRI metrics in the neonatal cortex. Specifically, regional variation in cortical morphometry and microstructure reflects differences in developmental maturity and tissue composition across cortical areas, indexed by the differential timing of gene expression across multiple cell types in the fetal cortex. Having established this relationship, we found that interruption to gestation through preterm birth resulted in significant disruptions to MRI-based measures of cortical development by the time of full-term birth. Further, the effects of preterm birth are temporally and spatially coincident to developmental processes involving cortical glial cell populations. This work provides an experimental framework to link molecular developmental mechanisms to macroscopic measures of cortical anatomy in early life, demonstrating the relationship between fetal gene expression and neonatal brain development and highlighting the specific impact of early exposure to the extrauterine environment due to preterm birth.

Using advanced MRI acquired close to the time of birth in a large, healthy neonatal population, we mapped multiple measures of regional cortical morphometry onto a single mode of variation, defining a principal axis of the neonatal cortex. Ordering of cortical regions along this axis separated lower order sensory and motor regions from higher-order regions including parietal, frontal and superior temporal cortex situated at opposite ends. The shared spatial ordering of cortical properties is a common organisational feature of the mammalian brain,[1,2,60,61] reflected in regional variations in cell populations,[61] gene expression,[22,62] and connectivity[60] as well as MRI-based measures of functional topography[5] and cortical morphometry in both adults[63] and infants.[64] The optimal mapping of cortical properties onto one or two lower dimensions remains an area of active research,[1] however, studies have demonstrated that variation along one axis, or gradient, is largely reflected by concerted changes in others,[2,6,65] suggesting that lower-order representations of cortical organisation largely capture shared views of latent neurobiological variation. An important benefit of this framework is the reduction of multiple metrics into a single measure per subject. In our case, this takes advantage of the inherent redundancy across multiple structural and diffusion MRI measures of the same cortical regions, producing a latent representation of cortical structure across scales. Here, we applied a simple linear mapping, arranging cortical regions along a single axis using PCA. This was sufficient to explain a significant proportion of variation in regional MRI-based metrics, with areas with similar cortical profiles clustering together along the principal axis. Based on an observed differential expression of genes associated with specific progenitor cell populations, order along this axis was correlated with spatial gene expression patterns that reflected a differential timing of cortical development across regions. Comparison to cell-specific gene expression profiles in late gestation suggested that MRI-based measures of cortical structure at birth correlated with gene expression by specific glial populations involving oligodendrocytes, microglia and endothelial cells.

This correlation potentially reflects a spatial variation in the developmental timing of processes associated with myelination, neuronal guidance and the continued maturation of the brain’s vascular networks at around the time of birth.[66–70]

The advent of modern transcriptomic technologies has enabled detailed analyses of the foundational molecular mechanisms underpinning corticogenesis in the human fetal brain.[16,52,53] Resolved to the level of individual cells, recent studies have performed systematic explorations of gene expression dynamics across cell-cycle progression, migration and differentiation of several major cell types in the fetal brain.[51] Combined with regional expression levels of bulk tissue mRNA measured across multiple cortical areas, this allows the spatio-temporal mapping of cell-specific gene expression profiles in the developing brain.[52] Here, we used a development atlas of gene expression, measured across 11 cortical regions from 12 to 37 post-conceptional weeks in 16 separate brain specimens.[18] This data resource provides unparalleled access to both the spatial and temporal dynamics of developmental mechanisms ongoing in the cortex during gestation. We found that a number of genes vary across cortical areas in line with a principal imaging axis. In particular, we found that genes with relatively higher expression in higher order regions during gestation were associated with developmentally earlier processes including neuronal differentiation and migration and were predominantly expressed by intermediate precursor cells and early-maturing inhibitory neurons. Using an alternative approach in four mid-gestation brain samples (aged 16-21 pcw), Miller et al. identified a generally rostro-caudal gradient of gene expression progressing along the contours of the developing brain and anchored in frontal and temporal cortex.[16] While some overlap was evident: 72/85 (85%) of frontally-enriched genes that were included in both studies were also positively correlated with the imaging axis, this indicates that variation along PC1 may reflect a combination of multiple overlapping intrinsic hierarchies or cellular gradients underlying cortical development.[18,61,62] Using a machine-learning approach designed to accommodate the large number of genes assayed, and validated in an independent sample, we established that the maturation of a given tissue sample could be accurately determined based on temporally-evolving profiles of gene expression. This approach takes advantage of the degree of variation in gene expression over development. Temporal variability in expression is present across most protein-coding genes and over 95% of genes that are differentially-expressed across cortical regions, are also differentially-expressed across gestation, with age explaining a large proportion of variance in gene expression.[18,19] Using the relative advancement or delay in predicted age across regional tissue samples, we observed a correlation between emerging differences in areal gene expression and cortical structure at birth, suggesting an interaction between the relative rate of development across regions and length of gestation. This was most notable in the protracted developmental trajectory of the visual cortex in midgestation, as noted elsewhere.[16] Overall, our results lend evidential support to the presence of heterogeneous corticogenic timing over gestation.[71,72]

Based on these observations, we hypothesised that interruption to gestation would lead to cortical disruptions along the principal cortical axis, reflecting a deleterious interaction with genetically-determined developmental programs ongoing in the cortex in the latter stages of gestation. To test this, we compared cortical development in healthy newborns to a cohort of preterm-born infants scanned at the time of normal birth. In line with previous observations,[28,43,73] we found significant differences across most cortical metrics of macro- and microstructure in the preterm brain. The magnitude of differences between cohorts aligned with the principal imaging axis suggesting a differential impact of perinatal adversity on cortical development that is potentially encoded by a selective vulnerability across regions due to differential maturation rates. Adverse intrauterine environments can result in altered patterns of fetal gene expression and brain development [74–79] and we demonstrate overlapping genetic associations between alterations in preterm cortical structure and differential expression in an experimental model of fetal hypoxic-ischaemic brain injury.[57] However, the antecedents and impacts of preterm birth on brain development are multifactorial and we remain cautious on speculating about the causal mechanisms that may underlie the relationships observed in this study without further empirical evidence.

The largest effect was observed in the myelin-sensitive T1w/T2w contrast. In adults, regional variation in cortical T1w/T2w contrast is high correlated with quantitative MRI-measures of intracortical myelin and histological maps of cytoarchitecture.[80] Myelination in the neonatal cortex is minimal, however T1w and T2w signal vary as a function of position in the neonatal cortex and the transcortical pattern of T1w/T2w ratio observed in this study mirrors closely that reported in older cohorts, with high values predominant in primary sensory regions.[80] In addition, we find that genes with expression correlated to T1w/T2w contrast are enriched for genes expressed by glial cells, including microglia and oligodendrocytes, across the second half of gestation. This mirrors earlier reports, based on microarray data, of correlations between neonatal imaging phenotypes and glial gene expression during gestation.[81] Using a time-resolved analysis, we found several molecular pathways involving genes with spatial and temporal correlation to the potential timing of preterm birth. This method leveraged nonlinear models fit using the full prenatal sample, allowing the discrete mapping of varying gene expression associations across the mid- to late-fetal period. We identify co-expressed networks of genes expressed by microglia and oligodendrocytes in late gestation and associated with Rho-GTPase signalling pathways, critical for neuronal migration [82] and involved in oligodendrocyte maturation and myelination [83,84]; cytokine signalling and inflammatory response pathways involving NF-κB1 and associated with microglial activation after hypoxic insult;[85] the MAPK/ERK signaling pathway, associated with neuronal and oligodendrocyte proliferation, [86,87] as well as sphingolipid metabolic pathways and apoptotic pathways expressed in endothelial cells. These data provide supporting evidence to the important role of developmental glial populations in preterm brain injury.[88–90] We have previously identified risk alleles in preterm born infants in genes involved in lipid metabolism and microglial activation in the developing brain and associated with altered patterns of brain development by term-equivalent age [91,92]. In this study, we validate our observations in an experimental model of fetal hypoxic-ischaemic injury highlighting a differential expression of glial-expressed genes after early brain injury.[57] We found several genes identified in both human and mouse studies and associated with T1/T2 differences in the preterm brain that were differentially-expressed after early brain injury suggested potential deleterious effects on glial cell populations that could lead to disrupted neuronal migration and formation of neural circuitry in the preterm brain.[57,57,93-96] While we recognise the need for further experimental research to elucidate the link to alterations in cortical structure, our findings highlight potential pathways by which preterm birth can result in altered cortical development due to coincidental timing with corticogenic processes in the fetal cortex.

We note several limitations to our study. While we have taken care to validate our observations across several datasets, we are careful to avoid causal language to describe the associations presented. Further experimental evidence is required to fully understand how spatial gene expression gradients lead to alterations in MRI-based measures of cortical structure in healthy and preterm brains. Due to the nature of the data, we compare *postmortem* gene expression data from the fetal cortex with *in vivo* measurements of cortical development in healthy and preterm neonates at the end of gestation. We recognise this approach depends upon a number of assumptions including that gene expression patterns can be generalised across preterm and healthy cohorts and that MRI-based metrics acquired at a single timepoint act as a surrogate measure for ongoing associations between cortical structure and gene expression during gestation. We mitigate some of these risks by performing validation experiments in independent datasets and comparing our findings to experimental models of preterm brain injury. To measure contemporaneous associations in imaging and gene expression, other studies [50] have employed *postmortem* MRI of fetal brains to acquire data in age-matched samples but this approach comes with the additional challenges of imaging *post mortem* tissue. With advancements in fetal MRI, we anticipate that future research will focus on examining imaging-transcriptomic associations across corresponding timepoints through mid- to late-gestation using healthy fetal MRI and preterm infants scanned shortly after birth to better capture the temporal evolution of the reported associations.

While the fetal dataset made available through PsychENCODE represents an unprecedented window into spatiotemporal gene expression in the developing brain, the relatively coarse spatial sampling limits our ability to map fine-grained spatial variation, or boundaries between primary and secondary areas. Our analyses only begins at 12pcw, after early gene expression gradients have begun to impose areal differentiation on the developing brain.[9] Similarly, the bulk tissue samples analysed contained fetal transient structures including the marginal zone and subplate, that differ from the cortex in terms of spatial and temporal development.[97] Although we verified our findings in the cortical plate using microarrays from layer-specific dissection in two 21pcw donor brains, this analysis was limited to a single timepoint. We anticipate, that with the increasing availability of spatially resolved gene expression datasets, advances in fetal and neonatal imaging, as well as layer-specific imaging analysis [98], further exploration of this area will yield interesting insight into early cortical development. In addition, sexual dimorphism in the transient fetal structures of the brain have been reported.[99] While we felt the relatively small sample size precluded a direct assessment of sex differences in gene expression, we included sex as a factor in all models of gene expression over time, and in our analyses of cortical MRI measures.

To examine cell-specific gene expression, we performed a stratification of bulk tissue RNA using gene lists collated from several independent scRNA studies. This method assumes that areal differences in gene expression are due to differences in developmental timing represented by cellular differentiation as well as changes in the proportion of cell types in composite tissue. While we did not test directly whether the same cell types also show areal differences in gene expression, this was previously explored by Fan et al. [51] In 22-23pcw samples, they found that the dominant mode of variation across cells was cell type rather than regional location and that most areal differences were driven by significant variations in tissue composition, as well as differing patterns of maturation. The advent of high-resolution single-cell RNA maps [53] will hopefully lead to future studies where we can more directly test regional development of specific cell types, rather than broader cell classes.

Finally, in this study, we focused on cortical structure rather than function. Future research may explore spatial associations between fetal gene expression and brain function as measured by MRI. In adults, a number of recent analyses have compared MRI metrics to patterns of gene expression reporting significant associations between correlated gene expression and both structural and functional measures [25,100–102] As discussed above, concerted variation in cortical properties is a common feature across MRI modalities [103,104] and spatial gene associations may be difficult to disentangle across correlated metrics. However, cortical measures that correlate at a single time point may not develop in tandem and future exploration of temporal development of cortical structure and function in relation to gene expression will yield interesting future research directions in this area.

In conclusion, we show that noninvasive imaging of the cortical structure in the neonatal brain is sensitive to differential spatiotemporal patterns of gene expression during gestation. In addition, we find that disruption to this developmental programming by preterm birth is associated with significant cortical alterations that appear to reflect the selective vulnerability of developing glial populations in the developing cortex.

## Author contributions

Conceptualisation: GB, JS, ADE; Data acquisition and processing: DC, JH, LC-G, EH, AP,GB, JO’M, RD, DF, AM, DC, AS, JP-P, ECR; Data analysis: GB, JS. Writing: GB, JS, JO’M, RD, ECR, ADE; Funding acquisition: ADE, JVH, DR

## Data and code availability

Neuroimaging data are available from http://www.developingconnectome.org/second-data-release/. RNA-seq data is available from http://development.psychencode.org/. Accession codes for additional gene expression data: BrainCloud GSE30272; mouse data GSE89998; single-cell RNA data GSE103723. LMD microarray data accessed via brainspan.org/lcm. Python and R code to perform the main analyses in the paper is available from https://github.com/garedaba/baby-brains.

## Materials and Methods

### Subjects

Infants were recruited and imaged at the Evelina Newborn Imaging Centre, St Thomas’ Hospital, London, UK for the Developing Human Connectome Project (dHCP). The study was approved by the UK Health Research Authority (Research Ethics Committee reference number: 14/LO/1169) and written parental consent was obtained for all participants. Neuroimaging and basic demographic data from the dHCP are available to download from: http://www.developingconnectome.org/second-data-release/.

In total, 442 healthy, term-born infants (gestational age at birth > 37 weeks) scanned between February 2015 and November 2018 as part of the dHCP were included in this study. From this cohort, n=362 were successfully processed via the dHCP structural processing pipeline (see *Image processing* below) and included after quality control. Of these, diffusion data from n=296 was successfully processed using both DTI and NODDI pipelines (see *Image processing* below) A further four subjects were excluded following a final visual inspection due to cropped anatomical images. Of 107 preterm infants (gestational age at birth < 37 weeks) scanned at term-equivalent age during the same period, one was excluded due to incomplete demographic data, n=84 completed structural MRI processing and n=67 passed diffusion processing after quality control. A further n=3 were removed after final visual inspection.

After quality control and image processing, the final cohort comprised n=292 healthy term-born infants (54% male, mean [S.D] postmenstrual age at birth=39.96 [1.10] weeks, mean [S.D.] age at imaging=40.94 [1.56] weeks) and n=64 preterm infants scanned at term-equivalent age (59% male; born 32.00 [3.88] weeks and imaged at 40.57 [2.25] weeks).

### Magnetic Resonance Imaging

MRI was performed on a 3T Philips Achieva (Philips, Netherlands) using a dedicated neonatal imaging system including a neonatal 32 channel phased array head coil.[29] Infants were imaged without sedation. T1- and T2-weighted anatomical images were acquired alongside diffusion and resting state functional MRI (total acquisition time: 63 minutes).

Inversion-recovery T1-weighted and T2-weighted images were acquired in sagittal and axial orientations (in-plane resolution: 0.8 × 0.8mm^2^, slice thickness: 1.6mm with 0.8mm overlap) with TR=4795ms; TI=1740ms; TE=8.7ms; SENSE: 2.27 (axial) and 2.66 (sagittal) for T1-weighted images and TR=12000ms, TE=156ms; SENSE: 2.11 (axial), 2.60 (sagittal) for T2-weighted. T1- and T2- weighted image stacks were motion corrected and reconstructed using the multi-slice aligned sensitivity encoding method with integration into a 3D volume using a super-resolution scheme into 0.8 × 0.8 × 0.8mm resolution volumes.[105,106]

Diffusion MRI was acquired with a spherically-optimised set of directions over 4 b-shells (b=0s/mm^2^: 20 directions; b=400: 64 directions; b=1000: 88 directions; b=2600: 128 directions) with a multiband factor acceleration of 4, TR=3800ms; TE=90ms; SENSE: 1.2 and acquired resolution of 1.5mm × 1.5mm with 3mm slices (1.5mm overlap) reconstructed using an extended SENSE technique into 1.5 × 1.5 × 1.5mm volumes.[107,108]

### Image processing

T1- and T2-weighted images were processing using the dHCP structural pipeline (https://github.com/BioMedIA/dhcp-structural-pipeline).[28] Briefly, T2-weighted images were bias corrected (N4),[109], brain-extracted (BET)[110] and segmented into grey matter, white matter and cerebrospinal fluid using DRAW-EM.[111] Cortical surfaces of the right and left hemisphere were then extracted[112] and aligned to a population-specific cortical template[113] using spherical inflation and multimodal surface matching (MSM) with higher order constraints (https://github.com/ecr05/MSM_HOCR).[114,115] This method ensures that all surfaces across participants have one-to-one vertex correspondence with the dHCP neonatal template. For each subject, we extracted the following metrics: cortical thickness (corrected for cortical curvature) and T1w/T2w contrast (calculated using rigidly aligned T1-weighted images).

Diffusion-weighted images were preprocessed by first denoising [116] and removing Gibbs ringing artefacts,[117] followed by a slice-to-volume motion and distortion correction with a slice-level outlier rejection using a multi-shell spherical harmonic signal representation (SHARD).[118] Visual inspection of the 4D images ensured motion correction and outlier rejection was successful and that images of poor quality were excluded from further analysis.

We fit each subject’s diffusion data with both a diffusion tensor model, fitted to the b=1000s^2^/mm shell and implemented in MRtrix,[119] and the NODDI (Neurite Orientation Dispersion and Density Imaging) model [120]. For the diffusion data, NODDI was implemented with the NODDI MATLAB toolbox using the *invivopreterm* tissue type options with the default parameters of intrinsic diffusivity fixed to 1.7 × 10^−3^ mm^2^/s and the starting point for values considered as the fraction of intra-neurite space lowered to 0-0.3 (instead of 0-1 in the adult brain) to better fit higher water content in the newborn compared to the mature adult brain.[48,121]

From these models, we derived parametric maps of fractional anisotropy (FA) and mean diffusivity (MD) from DTI, as well as maps of orientation dispersion index (ODI), quantifying the angular variation of neurite orientation within a voxel and intra-cellular volume fraction (fICVF), indexing the tissue volume fraction restricted within neurites. Cortical diffusion maps were projected to the cortical surface after co-registration with the corresponding anatomical data.

Images were visually inspected after acquisition and after reconstruction, and following each processing pipeline. Any images that failed to successfully complete the processing pipelines or failed visual inspection at any stage were removed from further analysis. As an additional step, we quantified in-scanner movement and image quality using a summary metric of the total head translation, rotation and the ratio of detected outlier slices. These three metrics were combined into one aggregated quality assurance measure [118,122] This measure did not significantly differ between groups (term [mean +/− S.D] = 1.61 +/− 1.79, preterm = 1.55 +/− 1.03; t=0.24, p=0.81). Including QA as a covariate in our analyses of group differences didn’t impact our reported observations.

### Bulk tissue gene expression data

Preprocessed, bulk tissue cortical gene expression data were made available as part of the PsychENCODE project (available to download at: http://development.psychencode.org/).[18] Tissue was collected after obtaining parental or next of kin consent and with approval by the institutional review boards at the Yale University School of Medicine, the National Institutes of Health, and at each institution from which tissue specimens were obtained.

Tissue processing is detailed elsewhere. [18] In brief, mRNA data were available for post-mortem human brain tissue collected from n=41 specimens aged between 8 post-conceptional weeks (pcw) and 40 postnatal years. For each brain, regional dissection of up to 16 cerebral regions was performed, including 11 neocortical regions (dorsolateral frontal cortex, DLPFC; ventrolateral frontal cortex, VLPFC; orbitofrontal cortex, OFC; medial frontal cortex, MFC; primary motor cortex, M1; primary sensory cortex, S1; inferior parietal cortex, IPC; primary auditory cortex, A1C; superior temporal cortex, STC; inferior temporal cortex, ITC; primary visual cortex, V1), and five sub-cortical regions (hippocampus, amygdala, striatum, thalamus and cerebellar cortex). Detailed anatomical boundaries for each cortical region at each stage of development are provided elsewhere.[17,18] Regional tissue samples were subject to mRNA-sequencing using an Illumina Genome Analyzer IIx (Illumina, San Diego, CA) and mRNA-seq data processed using RSEQtools (v0.5).[123] Gene expression was measured as reads per kilobase of transcript per million mapped reads (RPKM). Conditional quantile normalisation was performed to remove GC-content bias and ComBat used to remove technical variance due to processing site (Yale or USC).[18,124,125]

In this study, we included RPKM data from neocortical samples of prenatal specimens aged 12 post-conceptional weeks onwards (n=16, age range = 12-37 pcw, mean [S.D.] age = 18.4 [7.7] pcw, 50% male, mean [S.D.] number of cortical regions sampled = 9.75 [1.6], mean [S.D.] post-mortem interval = 7.1 [12.6] hours, mean [S.D.] RNA integrity number[RIN][126] = 9.26 [0.73]). Prenatal specimens from the earliest developmental window (8-9 postconceptional weeks) were excluded as some cortical regions (e.g.: M1 and S1) were combined together to account for immature cortical anatomy.[17,18]

The prenatal gene expression data was initially filtered to only include protein-coding genes (NCBI GRCh38.p12, n=18,524 out of a possible 20,720). In order to restrict our analysis to focus on genes expressed in the developing cortex, we further filtered this list to only contain genes expressed by cells in the fetal cortex based on the composite list of prenatal cell markers from five independent single-cell RNA studies of the developing fetal cortex (see *‘Genetic markers of cell type’* below). This resulted in expression data from a final set of 5287 genes.

### Additional gene expression datasets

#### BrainCloud

Preprocessed microarray data from n=46 human prefrontal cortex tissue samples aged approximately 95 to 390 postconceptional-days (14 pcw to 4 months postnatal age) were downloaded from GEO (accession: GSE30272). Prior to analysis, individual gene expression was modelled using nonlinear splines. Surrogate Variable Analysis was performed to remove technical variation and batch effects (31 surrogate variables) while retaining variation due to age. For further details please see Colantuoni et al. [56] For each gene, expression was Z-transformed prior to modelling. In total, gene expression for 4986/5287 fetal gene markers was available.

#### Single cell RNA

Regional single cell RNA gene expression data was made available via GEO (accession: GSE103723).[51] Briefly, 4213 single cells were isolated from 20 anatomical regions of the 22 and 23pcw fetal cortex and subject to single cell RNA-sequencing. Normalised expression data in transcripts per million (TPM) were available for 96 cells per tissue sample. We selected data from single cells extracted from matching cortical regions to those described above and classified to one of 10 classes based on clusters identified in [51], as detailed *‘Genetic markers of cell type’* below (n=572 cells).

#### Laser microdissection (LMD) microarray data

LMD microarray data was accessed via the BrainSpan data portal (brainspan.org/lcm). This provides access to DNA microarray data from four midgestational brains, dissected into around 300 anatomical samples. Detailed information is provided elsewhere.[16] We performed a differential search to identify microarray probes with differential expression in the cortical plate of the DLPFC (regional ID: fCPdl), VLPFC (fCPvl) and IPC (pCPpv) compared to M1 (fCPm1) and S1 (fCPs1) measured in two 21pcw donor brains. Differential expression data for 23,000 probes was downloaded and corresponding data mapped to the set of preselected fetal marker genes for comparison.

#### Experimental mouse model of preterm brain injury

Gene expression levels in P1.5 mouse cortex was measured for control or ischaemic pups, where ischaemia was induced by maternal uterine artery occlusion at E16.5. Four mice in each group were included. [57] Data were made available via GEO (accession: GSE89998). Analysis was performed using *GEO2R* and the *limma* package. Expression data were first log2-transformed before fold-change was estimated across groups. Mouse genes were mapped to human homologs using Ensembl and matched to the list of human fetal gene markers.

### Cortical regions-of-interest

To facilitate comparison between developmental RNA and MRI data, we created a set of cortical regions-of-interest (ROI) labels corresponding to the anatomical dissections used for mRNA analysis and aligned to the dHCP imaging data.

Reference post-mortem MRI data were acquired as part of the Allen Institute BrainSpan Atlas of the Developing Human Brain. Details of tissue processing and MRI acquisition are available at: https://help.brain-map.org/download/attachments/3506181/BrainSpan_MR_DW_DT_Imaging.pdf. In brief, MRI was acquired at 3T (Siemens, Germany) in a post-mortem, whole-brain specimens aged 22 pcw. In addition, anatomical annotations corresponding to the regional dissections in Miller et al.,[16] Kang et al.[17] and Li et al.[18] were provided on a reconstructed cortical surface from a 19pcw prenatal specimen.[50] Cortical ROI data were available to download in VTK file format, separately for left and right cortical hemispheres (Figure S1).

To generate a set of dHCP-compatible cortical labels, we reconstructed the cortical surface of a 3T post-mortem MRI image from a 22pcw brain. First, manually creating a brain mask to remove non-brain tissue, then smoothing using a mean filter of 3mm width.We performed automated tissue segmentation on the smoothed image using the dHCP structural pipeline, manually correcting tissue segmentations on a slice-by-slice basis for accuracy prior to cortical surface reconstruction. Using dHCP tools, the fetal cortical surface was extracted and cortical labels manually transferred onto it based on the reference labels provided by Huang et al.[50] and anatomical descriptions in Li et al.[18] Finally, the fetal surface was inflated to a sphere and co-registered to the earliest timepoint (36 weeks gestational age) of the dHCP cortical surface atlas using multimodal surface matching (MSM).[113,115]

This resulted in a set of 11 cortical ROI, each associated with regional bulk tissue mRNA data sampled across gestation and co-registered with dHCP neuroimaging data to allow correspondent sampling of cortical imaging metrics in the neonatal brain (Fig S1).

### Cortical imaging metric analysis

For every subject, mean values of each imaging metric (thickness, T1w/T2w contrast, FA, MD, fICVF, ODI) were calculated within each cortical ROI. Metric values were averaged across hemispheres and outlier values identified and removed using a median absolute deviation (MAD) of > 3.5.

For all healthy term-born infants, regional metrics were Z-transformed and averaged across subjects to produce a group average region × metric matrix representing the relative variation of each imaging metric across cortical regions.

We projected the group average data onto two axes using Principal Component Analysis (PCA) via eigendecomposition of the data covariance matrix. This results in a set of *L* eigenvectors, *W*_*L*_, that map the original *n*×*p* data matrix, *X* onto a set of orthogonal axes as: *T*_*L*_ = *XW*_*L*_. As generally *L* < *p*, the truncated *n*×*L* matrix, *T*_*L*_, forms a low-dimensional representation of the original data. We can then project each subject’s region × metric matrix, *X*_*S*_, onto a common set of axes as 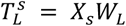, where 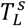 represents the *L* component scores for each subject, *s*.

All analysis was performed in Python (3.7.3) using Scipy (1.3.0)[127] and Scikit-Learn (0.21.2).[128]

### Modelling gene expression trajectories

For each gene, we modelled the relationship between gene expression and specimen age using mixed-effects models. Using bulk tissue RPKM data described above, each gene’s expression data were first Winsorised to set very small or large outlying values to the 5th and 95th centile values, respectively, to stabilise against extreme values before log2-transformation.

We compared two models, modelling regional gene expression as either a linear or nonlinear function of age with fixed effects of sex and RNA integrity number. We accounted for sample-specific variation by including in the model a random intercept for each specimen, such that:

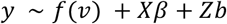

Where *f*(⋅) is a nonlinear function of predictor *v*, *X* is an *m*-observation × *p* design matrix modelling *p* linear, fixed effects and *Z* is an *m* ×(*n* ⋅ *r*) design matrix modelling *r* random effects across *n* specimens. In this case, age was included as either a nonlinear predictor, *f*(*v*), or as a fixed linear effect alongside sex and RIN. We specified a relatively smooth nonlinear function of age using a natural cubic spline with four knots evenly spaced across the age span. To estimate region-specific trajectories, we calculated a second nonlinear model, additionally including separate smooth functions for each cortical region. Models were compared using AIC and BIC.

We calculated age-corrected RPKM values for each gene in all cortical samples using the residuals of the best-fit nonlinear mixed model (Fig S3) to test the spatial association between gene expression and the principal imaging gradient using non-parametric correlation (Kendall’s *τ*).

Modelling was performed in R (3.6.1) using *nlme*[129] and *mgcv*[130] packages.

### Genetic markers of cell type

Genetic markers of cortical cell types were collated from five independent single-cell RNA studies of the fetal cortex.[18,51–54] Using single-cell RNA-seq, each study identified sets of genes differentially expressed across cell clusters or types. Cell types were independently defined in each study and a list of all cell types included in this study (n=87) are shown in Table S2. Where applicable, for a given cell type, differentially-expressed genes were included as cell type markers if they were found to be expressed in at least 50% all cells surveyed.[18,51,52] Across all five studies, each cell type was manually assigned to one of 11 cell classes based on text descriptions from each study (astrocyte, endothelial cell, microglia, neuron:excitatory, neuron:inhibitory, neuron:unclassified, oligodendrocyte, oligodendrocyte precursor cell [OPC], pericyte, intermediate progenitor cell, radial glia) and classified as either a precursor or mature cell type (Table S2). For each cell class, omnibus gene lists were created by collating identified gene markers for all cell types within a class. Unique gene lists were created by excluding any genes identified as a marker of more than one cell class.

### Cell type embedding

Using the region-specific, nonlinear model specified above, expression trajectories for every gene were estimated for each region at 50 evenly spaced points across the full observation window (12pcw - 37pcw). For each cell type identified in the fetal cortex (see above), expression trajectories for all cell-type gene markers were normalised to unit length, concatenated over regions and averaged to capture both temporal and spatial variation in average gene expression across cell types. Similarity between cell-type gene expression trajectories were then visualised by embedding into a two-dimensional space using Uniform Manifold and Approximation Projection (UMAP) based on Euclidean distance.[131]

### Enrichment analyses

We performed over-representation analysis (ORA) of each list of gene markers for each of 10 cell classes (excluding neuron:unclassified), calculating the hypergeometric statistic:

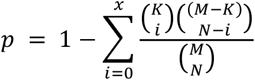

Where *p* is the probability of finding *x* or more genes from a cell-class-specific gene list *K* in a set of randomly selected genes, *N* drawn from a background set, *M*. We calculated enrichment ratios as the proportion of cell-class-specific genes in the gene list of interest, compared to the proportion in the full background set. The background gene set was defined as the full list of protein-coding genes included in the analysis (n=5287) unless otherwise specified. We corrected for multiple comparisons across cell classes using False Discovery Rate (FDR).

We additionally performed ORA for Gene Ontology terms using WebGestalt.[55]

### Weighted Gene Correlation Network Analysis

We used WGCNA [132] to identify co-expression modules within PC+ and PC- gene sets. We performed topology analysis using a gene × gene adjacency matrix constructed from the residualised log2-transformed RPKM data, after accounting for variance due to age, sex and sample effects (see *Modelling gene expression trajectories*, above). A soft threshold was chosen to approximate scale free topology in the adjacency matrix (PC+: power=5, r^2^=0.77; PC-: power=10, r^2^=0.78), [133] before transformation into a topological overlap matrix. Hierarchical clustering was used to assign genes to modules based on the dynamic tree-cutting method.[134] Analysis was performed in R (3.6.1) with the *WGCNA* package.[132]

### Predicting tissue maturity

We used gene expression over time to construct a predictive model of genetic maturity using support vector regression. To maximise coverage across the prenatal period, we included additional samples aged 8pcw up to 4 months of postnatal age (n=23 total). Using the n=120 regionally-varying genes (PC+ and PC-), we first calculated regional gene expression profiles, corrected for variance due to sex, RIN and specimen ID while retaining variance due to age, using nonlinear mixed-effects models. We then averaged gene expression across cortical regions in each specimen to create a specimen × gene (23 × 120) mean gene expression matrix, where each row represents the normalised log2(RPKM) of each gene for a given specimen, averaged across cortical regions.

In machine learning, kernels can be applied high-dimensional data sets to improve model fitting where *n*≪*p*. To calculate regional variation in genetic maturity, we implemented a leave-one-out (LOO) model using Support Vector Regression with a linear kernel (Scikit-Learn; regularisation parameter set to *C*=10.0) and modelling the association between specimen age (in post-conceptional days) and mean cortical gene expression data in 22 out of 23 specimens. We then used this model to predict age using the *regional* gene expression profiles of the remaining, left-out specimen, resulting in eleven age predictions, one per cortical region. We repeated this process, leaving out a different specimen each time.

In order to estimate a stable prediction of tissue maturity, we repeated the modelling using a bootstrapped selection of genes, repeating gene sampling with replacement 1000 times. We also repeated the modelling using all 5287 genes. We calculated the correlation between regional genetic maturity (averaged over 1000 bootstraps) and PC1 score for each specimen and tested the significance of this relationship by permuting mean gene expression profiles with respect to specimen age 5000 times during model training.

### Group comparison of cortical morphology

We compared regional cortical metrics in term and preterm cohorts using a linear mixed effects modelling approach. For each of six metrics, we modelled metric value as a combination of age, sex, regional PC1 score and birth group status (term or preterm). We included an interaction term for PC1 and birth status to test the hypothesis that preterm birth incurs differential effects across cortical regions in line with PC1. We also included subject ID as a random effect to account for correlated within-subject observations across regions. We fit nested models by Maximum Likelihood, comparing model fits with and without the inclusion of birth status using AIC and BIC (Table S5).

### Developmental gene enrichment

In order to test cell class enrichment over time, we split the preterm period (approximately 160 to 260 post-conceptional days) into 10 age windows. Using nonlinear gene expression trajectories, calculated across cortical regions (see *Modelling gene expression trajectories* above), we averaged modelled gene expression within each window for every cortical region. Then, in each window, we calculated the non-parametric association (Kendall’s *τ*) between gene expression and the mean difference between term and preterm groups in T1w/T2w contrast in each cortical region and recorded significantly associated genes (FDR-corrected at p<0.05). Finally, we performed cell-class enrichment (see ‘Enrichment Analyses’ above), in each of the 10, time-resolved gene sets.

### PPI networks

Protein-protein interactions networks were visualised in Cytoscape (3.7.2) using the StringDB protein query. Pathway enrichment of Reactome pathways was performed for subnetworks.

## References

1. Huntenburg JM, Bazin P-L, Margulies DS. Large-Scale Gradients in Human Cortical Organization. Trends in Cognitive Sciences. 2018;22: 21–31. doi:10.1016/j.tics.2017.11.002

2. Hilgetag CC, Goulas A. ‘Hierarchy’ in the organization of brain networks. Philosophical Transactions of the Royal Society B: Biological Sciences. 2020;375: 20190319. doi:10.1098/rstb.2019.0319

3. Raut RV, Snyder AZ, Raichle ME. Hierarchical dynamics as a macroscopic organizing principle of the human brain. PNAS. 2020 [cited 13 Aug 2020]. doi:10.1073/pnas.2003383117

4. Harris JA, Mihalas S, Hirokawa KE, Whitesell JD, Choi H, Bernard A, et al. Hierarchical organization of cortical and thalamic connectivity. Nature. 2019;575: 195–202. doi:10.1038/s41586-019-1716-z

5. Margulies DS, Ghosh SS, Goulas A, Falkiewicz M, Huntenburg JM, Langs G, et al. Situating the default-mode network along a principal gradient of macroscale cortical organization. PNAS. 2016;113: 12574–12579. doi:10.1073/pnas.1608282113

6. Burt JB, Demirtaş M, Eckner WJ, Navejar NM, Ji JL, Martin WJ, et al. Hierarchy of transcriptomic specialization across human cortex captured by structural neuroimaging topography. Nat Neurosci. 2018;21: 1251–1259. doi:10.1038/s41593-018-0195-0

7. Lein ES, Belgard TG, Hawrylycz M, Molnár Z. Transcriptomic Perspectives on Neocortical Structure, Development, Evolution, and Disease. Annu Rev Neurosci. 2017;40: 629–652. doi:10.1146/annurev-neuro-070815-013858

8. Silbereis JC, Pochareddy S, Zhu Y, Li M, Sestan N. The Cellular and Molecular Landscapes of the Developing Human Central Nervous System. Neuron. 2016;89: 248–268. doi:10.1016/j.neuron.2015.12.008

9. Bishop KM, Rubenstein JLR, O’Leary DDM. Distinct Actions of Emx1, Emx2, andPax6 in Regulating the Specification of Areas in the Developing Neocortex. J Neurosci. 2002;22: 7627–7638. doi:10.1523/JNEUROSCI.22-17-07627.2002

10. Kiecker C, Lumsden A. Hedgehog signaling from the ZLI regulates diencephalic regional identity. Nat Neurosci. 2004;7: 1242–1249. doi:10.1038/nn1338

11. Bystron I, Blakemore C, Rakic P. Development of the human cerebral cortex: Boulder Committee revisited. Nat Rev Neurosci. 2008;9: 110–122. doi:10.1038/nrn2252

12. Cadwell CR, Bhaduri A, Mostajo-Radji MA, Keefe MG, Nowakowski TJ. Development and Arealization of the Cerebral Cortex. Neuron. 2019;103: 980–1004. doi:10.1016/j.neuron.2019.07.009

13. Meyer G, Schaaps JP, Moreau L, Goffinet AM. Embryonic and Early Fetal Development of the Human Neocortex. J Neurosci. 2000;20: 1858–1868. doi:10.1523/JNEUROSCI.20-05-01858.2000

14. Kostovic I, Jovanov-Milosevic N. The development of cerebral connections during the first 20-45 weeks’ gestation. SeminFetalNeonatal Med. 2006;11: 415–422. doi:10.1016/j.siny.2006.07.001

15. Kostović I, Jovanov-Milošević N, Radoš M, Sedmak G, Benjak V, Kostović-Srzentić M, et al. Perinatal and early postnatal reorganization of the subplate and related cellular compartments in the human cerebral wall as revealed by histological and MRI approaches. Brain Struct Funct. 2014;219: 231–253. doi:10.1007/s00429-012-0496-0

16. Miller JA, Ding S-L, Sunkin SM, Smith KA, Ng L, Szafer A, et al. Transcriptional landscape of the prenatal human brain. Nature. 2014;508: 199–206. doi:10.1038/nature13185

17. Kang HJ, Kawasawa YI, Cheng F, Zhu Y, Xu X, Li M, et al. Spatio-temporal transcriptome of the human brain. Nature. 2011;478: 483–489. doi:10.1038/nature10523

18. Li M, Santpere G, Imamura Kawasawa Y, Evgrafov OV, Gulden FO, Pochareddy S, et al. Integrative functional genomic analysis of human brain development and neuropsychiatric risks. Science. 2018;362. doi:10.1126/science.aat7615

19. Pletikos M, Sousa AMM, Sedmak G, Meyer KA, Zhu Y, Cheng F, et al. Temporal Specification and Bilaterality of Human Neocortical Topographic Gene Expression. Neuron. 2014;81: 321–332. doi:10.1016/j.neuron.2013.11.018

20. Gandal MJ, Haney JR, Parikshak NN, Leppa V, Ramaswami G, Hartl C, et al. Shared molecular neuropathology across major psychiatric disorders parallels polygenic overlap. Science. 2018;359: 693–697. doi:10.1126/science.aad6469

21. Velmeshev D, Schirmer L, Jung D, Haeussler M, Perez Y, Mayer S, et al. Single-cell genomics identifies cell type-specific molecular changes in autism. Science. 2019;364: 685–689. doi:10.1126/science.aav8130

22. Hawrylycz MJ, Lein ES, Guillozet-Bongaarts AL, Shen EH, Ng L, Miller JA, et al. An anatomically comprehensive atlas of the adult human brain transcriptome. Nature. 2012;489: 391–399. doi:10.1038/nature11405

23. Wang D, Liu S, Warrell J, Won H, Shi X, Navarro FCP, et al. Comprehensive functional genomic resource and integrative model for the human brain. Science. 2018;362. doi:10.1126/science.aat8464

24. Fornito A, Arnatkevičiūtė A, Fulcher BD. Bridging the Gap between Connectome and Transcriptome. Trends Cogn Sci (Regul Ed). 2019;23: 34–50. doi:10.1016/j.tics.2018.10.005

25. Shin J, French L, Xu T, Leonard G, Perron M, Pike GB, et al. Cell-Specific Gene-Expression Profiles and Cortical Thickness in the Human Brain. Cereb Cortex. 2018;28: 3267–3277. doi:10.1093/cercor/bhx197

26. Richiardi J, Altmann A, Milazzo A-C, Chang C, Chakravarty MM, Banaschewski T, et al. Correlated gene expression supports synchronous activity in brain networks. Science. 2015;348: 1241–1244. doi:10.1126/science.1255905

27. Romero-Garcia R, Warrier V, Bullmore ET, Baron-Cohen S, Bethlehem RAI. Synaptic and transcriptionally downregulated genes are associated with cortical thickness differences in autism. Molecular Psychiatry. 2018; 1. doi:10.1038/s41380-018-0023-7

28. Makropoulos A, Robinson EC, Schuh A, Wright R, Fitzgibbon S, Bozek J, et al. The developing human connectome project: A minimal processing pipeline for neonatal cortical surface reconstruction. NeuroImage. 2018;173: 88–112. doi:10.1016/j.neuroimage.2018.01.054

29. Hughes EJ, Winchman T, Padormo F, Teixeira R, Wurie J, Sharma M, et al. A dedicated neonatal brain imaging system. Magnetic Resonance in Medicine. 2017;78: 794–804. doi:10.1002/mrm.26462

30. Knickmeyer RC, Gouttard S, Kang C, Evans D, Wilber K, Smith JK, et al. A Structural MRI Study of Human Brain Development from Birth to 2 Years. J Neurosci. 2008;28: 12176–12182. doi:10.1523/JNEUROSCI.3479-08.2008

31. Gilmore JH, Knickmeyer RC, Gao W. Imaging structural and functional brain development in early childhood. Nat Rev Neurosci. 2018;19: 123–137. doi:10.1038/nrn.2018.1

32. Kapellou O, Counsell SJ, Kennea N, Dyet L, Saeed N, Stark J, et al. Abnormal cortical development after premature birth shown by altered allometric scaling of brain growth. PLoS Med. 2006;3: e265. doi:10.1371/journal.pmed.0030265

33. Wang F, Lian C, Wu Z, Zhang H, Li T, Meng Y, et al. Developmental topography of cortical thickness during infancy. PNAS. 2019;116: 15855–15860. doi:10.1073/pnas.1821523116

34. Deoni SCL, Dean DC, O’Muircheartaigh J, Dirks H, Jerskey BA. Investigating white matter development in infancy and early childhood using myelin water faction and relaxation time mapping. Neuroimage. 2012;63: 1038–1053. doi:10.1016/j.neuroimage.2012.07.037

35. Counsell SJ, Maalouf EF, Fletcher AM, Duggan P, Battin M, Lewis HJ, et al. MR Imaging Assessment of Myelination in the Very Preterm Brain. AJNR Am J Neuroradiol. 2002;23: 872–881.

36. Doria V, Beckmann CF, Arichi T, Merchant N, Groppo M, Turkheimer FE, et al. Emergence of resting state networks in the preterm human brain. Proc Natl Acad Sci U S A. 2010;107: 20015–20020. doi:10.1073/pnas.1007921107

37. Fransson P, Åden U, Blennow M, Lagercrantz H. The Functional Architecture of the Infant Brain as Revealed by Resting-State fMRI. Cereb Cortex. 2011;21: 145–154. doi:10.1093/cercor/bhq071

38. Allievi AG, Arichi T, Tusor N, Kimpton J, Arulkumaran S, Counsell SJ, et al. Maturation of Sensori-Motor Functional Responses in the Preterm Brain. Cereb Cortex. 2016;26: 402–413. doi:10.1093/cercor/bhv203

39. Wilcox T, Haslup JA, Boas DA. Dissociation of processing of featural and spatiotemporal information in the infant cortex. Neuroimage. 2010;53: 1256–1263. doi:10.1016/j.neuroimage.2010.06.064

40. Cao M, He Y, Dai Z, Liao X, Jeon T, Ouyang M, et al. Early Development of Functional Network Segregation Revealed by Connectomic Analysis of the Preterm Human Brain. Cereb Cortex. 2017;27: 1949–1963. doi:10.1093/cercor/bhw038

41. Karen T, Morren G, Haensse D, Bauschatz AS, Bucher HU, Wolf M. Hemodynamic response to visual stimulation in newborn infants using functional near-infrared spectroscopy. Hum Brain Mapp. 2007;29: 453–460. doi:10.1002/hbm.20411

42. Ouyang M, Jeon T, Sotiras A, Peng Q, Mishra V, Halovanic C, et al. Differential cortical microstructural maturation in the preterm human brain with diffusion kurtosis and tensor imaging. PNAS. 2019;116: 4681–4688. doi:10.1073/pnas.1812156116

43. Ball G, Srinivasan L, Aljabar P, Counsell SJ, Durighel G, Hajnal JV, et al. Development of cortical microstructure in the preterm human brain. Proc Natl Acad Sci USA. 2013;110: 9541–9546. doi:10.1073/pnas.1301652110

44. Batalle D, O’Muircheartaigh J, Makropoulos A, Kelly CJ, Dimitrova R, Hughes EJ, et al. Different patterns of cortical maturation before and after 38 weeks gestational age demonstrated by diffusion MRI in vivo. NeuroImage. 2019;185: 764–775. doi:10.1016/j.neuroimage.2018.05.046

45. Yu Q, Ouyang A, Chalak L, Jeon T, Chia J, Mishra V, et al. Structural Development of Human Fetal and Preterm Brain Cortical Plate Based on Population-Averaged Templates. Cereb Cortex. 2016;26: 4381–4391. doi:10.1093/cercor/bhv201

46. Ball G, Boardman JP, Rueckert D, Aljabar P, Arichi T, Merchant N, et al. The effect of preterm birth on thalamic and cortical development. Cereb Cortex. 2012;22: 1016–1024. doi:10.1093/cercor/bhr176

47. Bouyssi-Kobar M, Brossard-Racine M, Jacobs M, Murnick J, Chang T, Limperopoulos C. Regional microstructural organization of the cerebral cortex is affected by preterm birth. NeuroImage: Clinical. 2018;18: 871–880. doi:10.1016/j.nicl.2018.03.020

48. Eaton-Rosen Z, Melbourne A, Orasanu E, Cardoso MJ, Modat M, Bainbridge A, et al. Longitudinal measurement of the developing grey matter in preterm subjects using multi-modal MRI. Neuroimage. 2015;111: 580–589. doi:10.1016/j.neuroimage.2015.02.010

49. Ouyang M, Dubois J, Yu Q, Mukherjee P, Huang H. Delineation of early brain development from fetuses to infants with diffusion MRI and beyond. Neuroimage. 2019;185: 836–850. doi:10.1016/j.neuroimage.2018.04.017

50. Huang H, Jeon T, Sedmak G, Pletikos M, Vasung L, Xu X, et al. Coupling Diffusion Imaging with Histological and Gene Expression Analysis to Examine the Dynamics of Cortical Areas across the Fetal Period of Human Brain Development. Cereb Cortex. 2013;23: 2620–2631. doi:10.1093/cercor/bhs241

51. Fan X, Dong J, Zhong S, Wei Y, Wu Q, Yan L, et al. Spatial transcriptomic survey of human embryonic cerebral cortex by single-cell RNA-seq analysis. Cell Res. 2018;28: 730–745. doi:10.1038/s41422-018-0053-3

52. Nowakowski TJ, Bhaduri A, Pollen AA, Alvarado B, Mostajo-Radji MA, Di Lullo E, et al. Spatiotemporal gene expression trajectories reveal developmental hierarchies of the human cortex. Science. 2017;358: 1318–1323. doi:10.1126/science.aap8809

53. Polioudakis D, de la Torre-Ubieta L, Langerman J, Elkins AG, Shi X, Stein JL, et al. A Single-Cell Transcriptomic Atlas of Human Neocortical Development during Mid-gestation. Neuron. 2019;103: 785–801.e8. doi:10.1016/j.neuron.2019.06.011

54. Pollen AA, Nowakowski TJ, Shuga J, Wang X, Leyrat AA, Lui JH, et al. Low-coverage single-cell mRNA sequencing reveals cellular heterogeneity and activated signaling pathways in developing cerebral cortex. Nat Biotechnol. 2014;32: 1053–1058. doi:10.1038/nbt.2967

55. Wang J, Vasaikar S, Shi Z, Greer M, Zhang B. WebGestalt 2017: a more comprehensive, powerful, flexible and interactive gene set enrichment analysis toolkit. Nucleic Acids Res. 2017;45: W130–W137. doi:10.1093/nar/gkx356

56. Colantuoni C, Lipska BK, Ye T, Hyde TM, Tao R, Leek JT, et al. Temporal dynamics and genetic control of transcription in the human prefrontal cortex. Nature. 2011;478: 519–523. doi:10.1038/nature10524

57. Kubo K-I, Deguchi K, Nagai T, Ito Y, Yoshida K, Endo T, et al. Association of impaired neuronal migration with cognitive deficits in extremely preterm infants. JCI Insight. 2017;2. doi:10.1172/jci.insight.88609

58. Szklarczyk D, Gable AL, Lyon D, Junge A, Wyder S, Huerta-Cepas J, et al. STRING v11: protein-protein association networks with increased coverage, supporting functional discovery in genome-wide experimental datasets. Nucleic Acids Res. 2019;47: D607–D613. doi:10.1093/nar/gky1131

59. Fabregat A, Jupe S, Matthews L, Sidiropoulos K, Gillespie M, Garapati P, et al. The Reactome Pathway Knowledgebase. Nucleic Acids Res. 2018;46: D649–D655. doi:10.1093/nar/gkx1132

60. Felleman DJ, Van Essen DC. Distributed hierarchical processing in the primate cerebral cortex. CerebCortex. 1991;1: 1–47.

61. Cahalane DJ, Charvet CJ, Finlay BL. Systematic, balancing gradients in neuron density and number across the primate isocortex. Front Neuroanat. 2012;6. doi:10.3389/fnana.2012.00028

62. Gomez J, Zhen Z, Weiner KS. Human visual cortex is organized along two genetically opposed hierarchical gradients with unique developmental and evolutionary origins. PLOS Biology. 2019;17: e3000362. doi:10.1371/journal.pbio.3000362

63. Wagstyl K, Ronan L, Goodyer IM, Fletcher PC. Cortical thickness gradients in structural hierarchies. Neuroimage. 2015;111: 241–250. doi:10.1016/j.neuroimage.2015.02.036

64. Larivière S, Vos de Wael R, Hong S-J, Paquola C, Tavakol S, Lowe AJ, et al. Multiscale Structure-Function Gradients in the Neonatal Connectome. Cereb Cortex. 2019 [cited 29 Nov 2019]. doi:10.1093/cercor/bhz069

65. Huntenburg JM, Bazin P-L, Goulas A, Tardif CL, Villringer A, Margulies DS. A Systematic Relationship Between Functional Connectivity and Intracortical Myelin in the Human Cerebral Cortex. Cereb Cortex. 2017;27: 981–997. doi:10.1093/cercor/bhx030

66. Fonta C, Imbert M. Vascularization in the Primate Visual Cortex during Development. Cereb Cortex. 2002;12: 199–211. doi:10.1093/cercor/12.2.199

67. Travis K, Ford K, Jacobs B. Regional dendritic variation in neonatal human cortex: a quantitative Golgi study. Dev Neurosci. 2005;27: 277–287. doi:10.1159/000086707

68. Jakovcevski I, Zecevic N. Sequence of oligodendrocyte development in the human fetal telencephalon. Glia. 2005;49: 480–491. doi:10.1002/glia.20134

69. Verney C, Monier A, Fallet-Bianco C, Gressens P. Early microglial colonization of the human forebrain and possible involvement in periventricular white-matter injury of preterm infants. J Anat. 2010;217: 436–448. doi:10.1111/j.1469-7580.2010.01245.x

70. Reemst K, Noctor SC, Lucassen PJ, Hol EM. The Indispensable Roles of Microglia and Astrocytes during Brain Development. Front Hum Neurosci. 2016;10. doi:10.3389/fnhum.2016.00566

71. Huttenlocher PR, Dabholkar AS. Regional differences in synaptogenesis in human cerebral cortex. Journal of Comparative Neurology. 1997;387: 167–178. doi:10.1002/(SICI)1096-9861(19971020)387:2<167::AID-CNE1>3.0.CO;2-Z

72. Guillery RW. Is postnatal neocortical maturation hierarchical? Trends in Neurosciences. 2005;28: 512–517. doi:10.1016/j.tins.2005.08.006

73. Dubois J, Benders M, Cachia A, Lazeyras F, Ha-Vinh Leuchter R, Sizonenko SV, et al. Mapping the early cortical folding process in the preterm newborn brain. CerebCortex. 2008;18: 1444–1454. doi:10.1093/cercor/bhm180

74. Garbett KA, Hsiao EY, Kálmán S, Patterson PH, Mirnics K. Effects of maternal immune activation on gene expression patterns in the fetal brain. Transl Psychiatry. 2012;2: e98. doi:10.1038/tp.2012.24

75. Tilley SK, Joseph RM, Kuban KCK, Dammann OU, O’Shea TM, Fry RC. Genomic biomarkers of prenatal intrauterine inflammation in umbilical cord tissue predict later life neurological outcomes. PLoS One. 2017;12. doi:10.1371/journal.pone.0176953

76. Edlow AG, Guedj F, Sverdlov D, Pennings JLA, Bianchi DW. Significant Effects of Maternal Diet During Pregnancy on the Murine Fetal Brain Transcriptome and Offspring Behavior. Front Neurosci. 2019;13. doi:10.3389/fnins.2019.01335

77. Miguel PM, Pereira LO, Silveira PP, Meaney MJ. Early environmental influences on the development of children’s brain structure and function. Developmental Medicine & Child Neurology. 2019;61: 1127–1133. doi:10.1111/dmcn.14182

78. Schneider C, Krischke G, Rascher W, Gassmann M, Trollmann R. Systemic hypoxia differentially affects neurogenesis during early mouse brain maturation. Brain Dev. 2012;34: 261–273. doi:10.1016/j.braindev.2011.07.006

79. Schwartz ML, Vaccarino F, Chacon M, Yan WL, Ment LR, Stewart WB. Chronic neonatal hypoxia leads to long term decreases in the volume and cell number of the rat cerebral cortex. Semin Perinatol. 2004;28: 379–388. doi:10.1053/j.semperi.2004.10.009

80. Glasser MF, Van Essen DC. Mapping Human Cortical Areas In Vivo Based on Myelin Content as Revealed by T1- and T2-Weighted MRI. J Neurosci. 2011;31: 11597–11616. doi:10.1523/JNEUROSCI.2180-11.2011

81. Pecheva D, Lee A, Poh JS, Chong Y-S, Shek LP, Gluckman PD, et al. Neural Transcription Correlates of Multimodal Cortical Phenotypes during Development. Cereb Cortex. [cited 15 Jan 2020]. doi:10.1093/cercor/bhz271

82. Govek E-E, Hatten ME, Van Aelst L. The Role of Rho GTPase Proteins in CNS Neuronal Migration. Dev Neurobiol. 2011;71: 528–553. doi:10.1002/dneu.20850

83. Barres BA, Raff MC, Gaese F, Bartke I, Dechant G, Barde YA. A crucial role for neurotrophin-3 in oligodendrocyte development. Nature. 1994;367: 371–375. doi:10.1038/367371a0

84. Feltri ML, Suter U, Relvas JB. The function of RhoGTPases in axon ensheathment and myelination. Glia. 2008;56: 1508–1517. doi:10.1002/glia.20752

85. Yao L, Kan EM, Kaur C, Dheen ST, Hao A, Lu J, et al. Notch-1 Signaling Regulates Microglia Activation via NF-κB Pathway after Hypoxic Exposure In Vivo and In Vitro. PLOS ONE. 2013;8: e78439. doi:10.1371/journal.pone.0078439

86. Ishii A, Furusho M, Bansal R. Sustained Activation of ERK1/2 MAPK in Oligodendrocytes and Schwann Cells Enhances Myelin Growth and Stimulates Oligodendrocyte Progenitor Expansion. J Neurosci. 2013;33: 175–186. doi:10.1523/JNEUROSCI.4403-12.2013

87. Samuels IS, Karlo JC, Faruzzi AN, Pickering K, Herrup K, Sweatt JD, et al. Deletion of ERK2 Mitogen-Activated Protein Kinase Identifies Its Key Roles in Cortical Neurogenesis and Cognitive Function. J Neurosci. 2008;28: 6983–6995. doi:10.1523/JNEUROSCI.0679-08.2008

88. Dommergues M-A, Plaisant F, Verney C, Gressens P. Early microglial activation following neonatal excitotoxic brain damage in mice: a potential target for neuroprotection. Neuroscience. 2003;121: 619–628. doi:10.1016/s0306-4522(03)00558-x

89. Back SA, Luo NL, Borenstein NS, Levine JM, Volpe JJ, Kinney HC. Late oligodendrocyte progenitors coincide with the developmental window of vulnerability for human perinatal white matter injury. JNeurosci. 2001;21: 1302–1312.

90. Baburamani AA, Supramaniam VG, Hagberg H, Mallard C. Microglia toxicity in preterm brain injury. Reproductive Toxicology. 2014;48: 106–112. doi:10.1016/j.reprotox.2014.04.002

91. Krishnan ML, Wang Z, Aljabar P, Ball G, Mirza G, Saxena A, et al. Machine learning shows association between genetic variability in PPARG and cerebral connectivity in preterm infants. PNAS. 2017;114: 13744–13749. doi:10.1073/pnas.1704907114

92. Van Steenwinckel J, Schang A-L, Krishnan ML, Degos V, Delahaye-Duriez A, Bokobza C, et al. Decreased microglial Wnt/β-catenin signalling drives microglial pro-inflammatory activation in the developing brain. Brain. 2019;142: 3806–3833. doi:10.1093/brain/awz319

93. Squarzoni P, Oller G, Hoeffel G, Pont-Lezica L, Rostaing P, Low D, et al. Microglia Modulate Wiring of the Embryonic Forebrain. Cell Reports. 2014;8: 1271–1279. doi:10.1016/j.celrep.2014.07.042

94. Stolp HB, Fleiss B, Arai Y, Supramaniam V, Vontell R, Birtles S, et al. Interneuron Development Is Disrupted in Preterm Brains With Diffuse White Matter Injury: Observations in Mouse and Human. Front Physiol. 2019;10. doi:10.3389/fphys.2019.00955

95. Fleiss B, Gressens P, Stolp HB. Cortical Gray Matter Injury in Encephalopathy of Prematurity: Link to Neurodevelopmental Disorders. Front Neurol. 2020;11: 575. doi:10.3389/fneur.2020.00575

96. Moretti R, Pansiot J, Bettati D, Strazielle N, Ghersi-Egea J-F, Damante G, et al. Blood-brain barrier dysfunction in disorders of the developing brain. Front Neurosci. 2015;9. doi:10.3389/fnins.2015.00040

97. Vasung L, Rollins CK, Velasco-Annis C, Yun HJ, Zhang J, Warfield SK, et al. Spatiotemporal Differences in the Regional Cortical Plate and Subplate Volume Growth during Fetal Development. Cereb Cortex. 2020;30: 4438–4453. doi:10.1093/cercor/bhaa033

98. Wagstyl K, Larocque S, Cucurull G, Lepage C, Cohen JP, Bludau S, et al. BigBrain 3D atlas of cortical layers: Cortical and laminar thickness gradients diverge in sensory and motor cortices. PLOS Biology. 2020;18: e3000678. doi:10.1371/journal.pbio.3000678

99. Vasung L, Rollins CK, Yun HJ, Velasco-Annis C, Zhang J, Wagstyl K, et al. Quantitative In vivo MRI Assessment of Structural Asymmetries and Sexual Dimorphism of Transient Fetal Compartments in the Human Brain. Cereb Cortex. 2020;30: 1752–1767. doi:10.1093/cercor/bhz200

100. Ritchie J, Pantazatos SP, French L. Transcriptomic characterization of MRI contrast with focus on the T1-w/T2-w ratio in the cerebral cortex. Neuroimage. 2018;174: 504–517. doi:10.1016/j.neuroimage.2018.03.027

101. Wang G-Z, Belgard TG, Mao D, Chen L, Berto S, Preuss TM, et al. Correspondence between resting state activity and brain gene expression. Neuron. 2015;88: 659–666. doi:10.1016/j.neuron.2015.10.022

102. Vértes PE, Rittman T, Whitaker KJ, Romero-Garcia R, Váša F, Kitzbichler MG, et al. Gene transcription profiles associated with inter-modular hubs and connection distance in human functional magnetic resonance imaging networks. Philos Trans R Soc Lond B Biol Sci. 2016;371. doi:10.1098/rstb.2015.0362

103. Baum GL, Cui Z, Roalf DR, Ciric R, Betzel RF, Larsen B, et al. Development of structure-function coupling in human brain networks during youth. PNAS. 2020;117: 771–778. doi:10.1073/pnas.1912034117

104. Vázquez-Rodríguez B, Suárez LE, Markello RD, Shafiei G, Paquola C, Hagmann P, et al. Gradients of structure-function tethering across neocortex. PNAS. 2019;116: 21219–21227. doi:10.1073/pnas.1903403116

105. Cordero-Grande L, Hughes EJ, Hutter J, Price AN, Hajnal JV. Three-dimensional motion corrected sensitivity encoding reconstruction for multi-shot multi-slice MRI: Application to neonatal brain imaging. Magn Reson Med. 2018;79: 1365–1376. doi:10.1002/mrm.26796

106. Kuklisova-Murgasova M, Quaghebeur G, Rutherford MA, Hajnal JV, Schnabel JA. Reconstruction of fetal brain MRI with intensity matching and complete outlier removal. Med Image Anal. 2012;16: 1550–1564. doi:10.1016/j.media.2012.07.004

107. Hutter J, Tournier JD, Price AN, Cordero-Grande L, Hughes EJ, Malik S, et al. Time-efficient and flexible design of optimized multishell HARDI diffusion. Magn Reson Med. 2018;79: 1276–1292. doi:10.1002/mrm.26765

108. Zhu K, Dougherty RF, Wu H, Middione MJ, Takahashi AM, Zhang T, et al. Hybrid-Space SENSE Reconstruction for Simultaneous Multi-Slice MRI. IEEE Trans Med Imaging. 2016;35: 1824–1836. doi:10.1109/TMI.2016.2531635

109. Tustison NJ, Avants BB, Cook PA, Zheng Y, Egan A, Yushkevich PA, et al. N4ITK: Improved N3 Bias Correction. IEEE Trans Med Imaging. 2010;29: 1310–1320. doi:10.1109/TMI.2010.2046908

110. Smith SM. Fast robust automated brain extraction. Human Brain Mapping. 2002;17: 143–155. doi:10.1002/hbm.10062

111. Makropoulos A, Gousias IS, Ledig C, Aljabar P, Serag A, Hajnal JV, et al. Automatic whole brain MRI segmentation of the developing neonatal brain. IEEE Trans Med Imaging. 2014;33: 1818–1831. doi:10.1109/TMI.2014.2322280

112. Schuh A, Makropoulos A, Wright R, Robinson EC, Tusor N, Steinweg J, et al. A deformable model for the reconstruction of the neonatal cortex. 2017 IEEE 14th International Symposium on Biomedical Imaging (ISBI 2017). 2017. pp. 800–803. doi:10.1109/ISBI.2017.7950639

113. Bozek J, Makropoulos A, Schuh A, Fitzgibbon S, Wright R, Glasser MF, et al. Construction of a neonatal cortical surface atlas using Multimodal Surface Matching in the Developing Human Connectome Project. Neuroimage. 2018;179: 11–29. doi:10.1016/j.neuroimage.2018.06.018

114. Robinson EC, Jbabdi S, Glasser MF, Andersson J, Burgess GC, Harms MP, et al. MSM: a new flexible framework for Multimodal Surface Matching. Neuroimage. 2014;100: 414–426. doi:10.1016/j.neuroimage.2014.05.069

115. Robinson EC, Garcia K, Glasser MF, Chen Z, Coalson TS, Makropoulos A, et al. Multimodal surface matching with higher-order smoothness constraints. Neuroimage. 2018;167: 453–465. doi:10.1016/j.neuroimage.2017.10.037

116. Veraart J, Fieremans E, Novikov DS. Diffusion MRI noise mapping using random matrix theory. Magnetic Resonance in Medicine. 2016;76: 1582–1593. doi:10.1002/mrm.26059

117. Kellner E, Dhital B, Kiselev VG, Reisert M. Gibbs-ringing artifact removal based on local subvoxel-shifts. Magnetic Resonance in Medicine. 2016;76: 1574–1581. doi:10.1002/mrm.26054

118. Christiaens D, Cordero-Grande L, Pietsch M, Hutter J, Price AN, Hughes EJ, et al. Scattered slice SHARD reconstruction for motion correction in multi-shell diffusion MRI of the neonatal brain. arXiv:190502996 [physics]. 2019 [cited 14 Nov 2019]. Available: http://arxiv.org/abs/1905.02996

119. Tournier J-D, Smith R, Raffelt D, Tabbara R, Dhollander T, Pietsch M, et al. MRtrix3: A fast, flexible and open software framework for medical image processing and visualisation. NeuroImage. 2019;202: 116137. doi:10.1016/j.neuroimage.2019.116137

120. Zhang H, Schneider T, Wheeler-Kingshott CA, Alexander DC. NODDI: practical in vivo neurite orientation dispersion and density imaging of the human brain. Neuroimage. 2012;61: 1000–1016. doi:10.1016/j.neuroimage.2012.03.072

121. Kunz N, Zhang H, Vasung L, O’Brien KR, Assaf Y, Lazeyras F, et al. Assessing white matter microstructure of the newborn with multi-shell diffusion MRI and biophysical compartment models. NeuroImage. 2014;96: 288–299. doi:10.1016/j.neuroimage.2014.03.057

122. Dimitrova R, Pietsch M, Christiaens D, Ciarrusta J, Wolfers T, Batalle D, et al. Heterogeneity in Brain Microstructural Development Following Preterm Birth. Cereb Cortex. 2020;30: 4800–4810. doi:10.1093/cercor/bhaa069

123. Habegger L, Sboner A, Gianoulis TA, Rozowsky J, Agarwal A, Snyder M, et al. RSEQtools: a modular framework to analyze RNA-Seq data using compact, anonymized data summaries. Bioinformatics. 2011;27: 281–283. doi:10.1093/bioinformatics/btq643

124. Hansen KD, Irizarry RA, Wu Z. Removing technical variability in RNA-seq data using conditional quantile normalization. Biostatistics. 2012;13: 204–216. doi:10.1093/biostatistics/kxr054

125. Johnson WE, Li C, Rabinovic A. Adjusting batch effects in microarray expression data using empirical Bayes methods. Biostatistics. 2007;8: 118–127. doi:10.1093/biostatistics/kxj037

126. Schroeder A, Mueller O, Stocker S, Salowsky R, Leiber M, Gassmann M, et al. The RIN: an RNA integrity number for assigning integrity values to RNA measurements. BMC Molecular Biology. 2006;7: 3. doi:10.1186/1471-2199-7-3

127. SciPy.org — SciPy.org. [cited 2 Oct 2019]. Available: https://www.scipy.org/

128. Pedregosa F, Varoquaux G, Gramfort A, Michel V, Thirion B, Grisel O, et al. Scikit-learn: Machine Learning in Python. Journal of Machine Learning Research. 2011;12: 2825–2830.

129. nlme: Nonlinear Mixed-Effects Models in nlme: Linear and Nonlinear Mixed Effects Models. [cited 2 Oct 2019]. Available: https://rdrr.io/cran/nlme/man/nlme.html

130. Wood S. Generalized Additive Models: An Introduction with R. 2nd ed. CRC Press; 2017. Available: https://www.crcpress.com/Generalized-Additive-Models-An-Introduction-with-R-Second-Edition/Wood/p/book/9781498728331

131. McInnes L, Healy J, Melville J. UMAP: Uniform Manifold Approximation and Projection for Dimension Reduction. arXiv:180203426 [cs, stat]. 2018 [cited 2 Oct 2019]. Available: http://arxiv.org/abs/1802.03426

132. Langfelder P, Horvath S. WGCNA: an R package for weighted correlation network analysis. BMC Bioinformatics. 2008;9: 559. doi:10.1186/1471-2105-9-559

133. Zhang B, Horvath S. A general framework for weighted gene co-expression network analysis. Stat Appl Genet Mol Biol. 2005;4: Article17. doi:10.2202/1544-6115.1128

134. Langfelder P, Zhang B, Horvath S. Defining clusters from a hierarchical cluster tree: the Dynamic Tree Cut package for R. Bioinformatics. 2008;24: 719–720. doi:10.1093/bioinformatics/btm563

